# MLT-11 is necessary for *C. elegans* embryogenesis and conserved sequences play distinct roles in cuticle structure

**DOI:** 10.1101/2025.11.14.688552

**Authors:** James Matthew Ragle, Ariela Turzo, Anton Jackson, An A. Vo, Vivian T. Pham, Keya Daly, John C. Clancy, Max T. Levenson, Alex D. Lee, Jordan D. Ward

## Abstract

Apical extracellular matrices (aECMs) are associated with many epithelia and many form a protective layer against biotic and abiotic threats in the environment. Despite their importance, we lack a deep understanding of their structure and dynamics in development and disease. *C. elegans* molting offers a powerful entry point to understanding developmentally programmed aECM remodeling. Here, we show that the poorly characterized putative protease inhibitor gene, *mlt-11*, is directly regulated by the NHR-23 transcription factor. We identify key *cis-*regulatory elements required for robust *mlt-11* expression. An internal MLT-11::mNeonGreen translational fusion transiently localized to the aECM in the cuticle and embryo. MLT-11::mNeonGreen was also detected in lining openings to the exterior (vulva, rectum, mouth). *mlt-11* is necessary to pattern all layers of the adult cuticle, and reduction of MLT-11 levels disrupted the barrier function of the cuticle. Deletion of conserved Kunitz protease inhibitor domains or intervening sequences produced a range of defects including either left or right rollers, and small separations of the cuticle along the length of the animal (microblisters). MLT-11 is processed into at least two fragments and internal and C-terminal mNeonGreen knock-ins display distinct localization patterns. Predicted *mlt-11* null mutations caused fully penetrant embryonic lethality and elongation defects. Together, this work suggests that MLT-11 localizes similarly to pre-cuticle components and conserved sequences play distinct roles in promoting proper assembly of the aECM.

## INTRODUCTION

Specialized extracellular matrices cover the apical surface of most epithelial cells and form the skin in almost all animals (Bonnans et al. 2014; Li Zheng et al. 2020; Diller and Tabor 2022). These apical extracellular matrices (aECMs) also line the lumen of internal tubular epithelia to form a protective layer against biotic and abiotic threats (Bonnans et al. 2014; Li Zheng et al. 2020; Diller and Tabor 2022). Despite their importance, understanding the structure and dynamics of aECM components in development and disease remains challenging.

*C. elegans* is a powerful model to study aECM structure and remodeling. Animals have a collagen-based aECM (cuticle) that may provide insight into mammalian skin biology dynamics (Page and Johnstone 2007; Sundaram and Pujol 2024). The components of the cuticle are secreted by hypodermal and seam cells and are assembled in distinct layers (Cox et al. 1981; Edgar et al. 1982; Page and Johnstone 2007). The cuticle secreted by the hypodermal syncytium form circumferential ridges called annuli separated from one another by furrows (Page and Johnstone 2007; Sundaram and Pujol 2024). In adults, an inner basal layer contains two fibrous sub-layers angled in opposite directions and an outer cortical layer (Edgar et al. 1982). A fluid-filled medial layer contains hollow, nano-scale struts built from the BLI-1, BLI-2, and BLI-6 collagens (Edgar et al. 1982; Adams et al. 2023). All layers are composed of extensively cross-linked collagens. The cortical layer also contains cuticlins, proteins which remain in the insoluble fraction after cuticle solubilization (Ristoratore et al. 1994). The cortical layer is covered by the epicuticle, a poorly understood structure that is thought to be a lipid bilayer covered by a glycoprotein rich surface coat (Blaxter 1993; Peixoto and De Souza 1995; Bada Juarez et al. 2019). There can be stage-specific variations in cuticle structure. Only adults have a medial layer, and both L1 and adult cuticles contain alae, lateral longitudinal ridges secreted by seam cells in the epidermis (Edgar et al. 1982; Katz et al. 2022). Dauer larvae, which are a specialized stress-resistant alternate L3 stage, have alae and a thicker cuticle than other stages and express unique collagens (Cox et al. 1980; Cox and Hirsh 1985).

During each larval stage animals must build a new aECM underneath the old one and separate the old aECM (apolysis) that is subsequently shed (ecdysis) (Lažetić and Fay 2017; Sundaram and Pujol 2024). A specialized, transient structure known as the pre-cuticle is thought to pattern the new cuticle and is then endocytosed (Cohen and Sundaram 2020). Pre-cuticle components include zona pellucida proteins, lipocalins, fibrillin, extracellular leucine-rich repeat proteins, and hedgehog-related proteins (Gill et al. 2016; Forman-Rubinsky et al. 2017; Cohen et al. 2019; Flatt et al. 2019; Cohen and Sundaram 2020; Serra and Sundaram 2021). The sheath is a similar structure in embryos which ensures embryonic membrane integrity and directs force during elongation (Priess and Hirsh 1986; Costa et al. 1997; Kelley et al. 2015; Vuong-Brender et al. 2017). The vulval aECM has recently been shown to be highly dynamic, and specialized aECMs also line the rectum, excretory system, mouth, and glial socket cells (Gill et al. 2016; Cohen et al. 2019; Cohen et al. 2020a; Kamal et al. 2022).

A major question is how is the new aECM constructed and the old one shed during molting? A poorly understood genetic oscillator is thought to coordinate the expression of cuticle components and processing enzymes (Hendriks et al. 2014; Meeuse et al. 2020). The molting cycle must also be tightly linked to developmental progression with homologs of circadian rhythm proteins (LIN-42/PER, NHR-23/ROR) playing important roles in both the molting (Kostrouchova et al. 1998; Kostrouchova et al. 2001; Monsalve et al. 2011; Spangler et al. 2025) and developmental timers (Jeon et al. 1999; Tennessen et al. 2006; Patel et al. 2022; Kinney et al. 2023). Study of NHR-23-regulated genes revealed an enrichment in predicted protease and protease inhibitors (Kouns et al. 2011; Johnson et al. 2023). Proteases are required for molting or ecdysis in both *C. elegans* and parasitic nematodes, presumably by promoting apolysis, though some are thought to function in collagen processing (Thacker et al. 1995; Davis et al. 2004; Hashmi et al. 2004; Frand et al. 2005; Stepek et al. 2010b; Kim et al. 2011; Stepek et al. 2011; Birnbaum et al. 2023). Proteins with Kunitz protease inhibitor domains have been implicated in molting through RNAi screening and have been suggested to suppress ecdysis (Frand et al. 2005; Stepek et al. 2010a; Lažetić and Fay 2017). MLT-11 is a predicted protease inhibitor in the Kunitz family and *mlt-11* RNAi causes molting defects (Frand et al. 2005). The aECM morphology mutant *rol-9* was recently mapped to the *mlt-11* gene, and the canonical *sc148* allele causes a semi-dominant right roller phenotype through an in-frame deletion in a conserved exon (Rich et al. 2022). *mlt-11* mRNA oscillates, peaking mid-molt (Hendriks et al. 2014; Meeuse et al. 2020) and its expression is regulated by NHR-23 (Frand et al. 2005), yet *mlt-11* remains poorly characterized.

Here we identify two NHR-23-regulated *cis-*regulatory elements important for *mlt-11* expression. Deletion of both elements or *mlt-11* RNAi caused developmental delay, motility defects, a defective cuticle barrier, and aberrant localization of the collagens BLI-1 and ROL-6. *mlt-11* RNAi consistently produced stronger phenotypes than *cis-*element deletion mutants. *mlt-11(RNAi)* animals displayed defective localization of aECM components in the basal, medial, and cortical layer. Structure-function analysis using precise deletions revealed a striking range of phenotypes. Depending on the sequence deleted, we observed embryonic lethality, right rollers, left rollers, or small detachments of the cuticle that we call microblisters (µBli). mNeonGreen::3xFLAG insertions at the C-terminus and internally produced distinct localization patterns and western blot analysis suggested that MLT-11 is post-translationally processed into at least two fragments. Internal MLT-11::mNG::3xFLAG displayed dynamic localization to the aECM in larvae and embryos and was endocytosed into lysosomes, similar to pre-cuticle factors. In contrast, a C-terminal MLT-11:mNG::3xFLAG exhibited weaker aECM localization and exhibited a more diffuse localization in embryos and the vulva. Embryonic lethality arises from a loss of cell junction integrity during elongation. This is the first work to define the *mlt-11* null phenotype and to confer that Kunitz domains and other conserved sequences may confer functional specificity.

## RESULTS

### *mlt-11* is expressed in embryonic and larval epidermal cells throughout development

To explore *mlt-11* expression we created a single copy *mlt-11p::mNeonGreen* (hereafter referred to as *mlt-11p::mNG*) promoter reporter using 2.8 kb of sequence upstream of the transcriptional start site. The reporter was expressed in embryos starting at the bean stage in posterior epithelial cells. Expression persisted through the 3-fold stage, spreading more anteriorly (Figure 1A). We also detected expression in epidermal cells in both larvae and adults (hyp7 syncytium and seam cells), similar to previous reports using an extrachromosomal array-based promoter reporter (Figure 1B) (Frand et al. 2005). We also detected expression in rectal and vulval cells (Figure 1B). A previous promoter reporter used 3 kb of sequence upstream but separated from the transcription start site (TSS) by about 2 kb of sequence (Frand et al. 2005). We created a single copy *mlt-11p::mNeonGreen::3xFLAG::PEST* promoter reporter containing this 3 kb of sequence as well as an additional reporter containing sequence spanning from the TSS to 5.3 kb upstream (Figure1C). Similar expression timing and intensity was seen in these two reporters compared to our original 2.8 kb promoter reporter (Figure1C), suggesting that both shorter promoter reporters captured key *cis-*regulatory elements.

**Figure 1.**
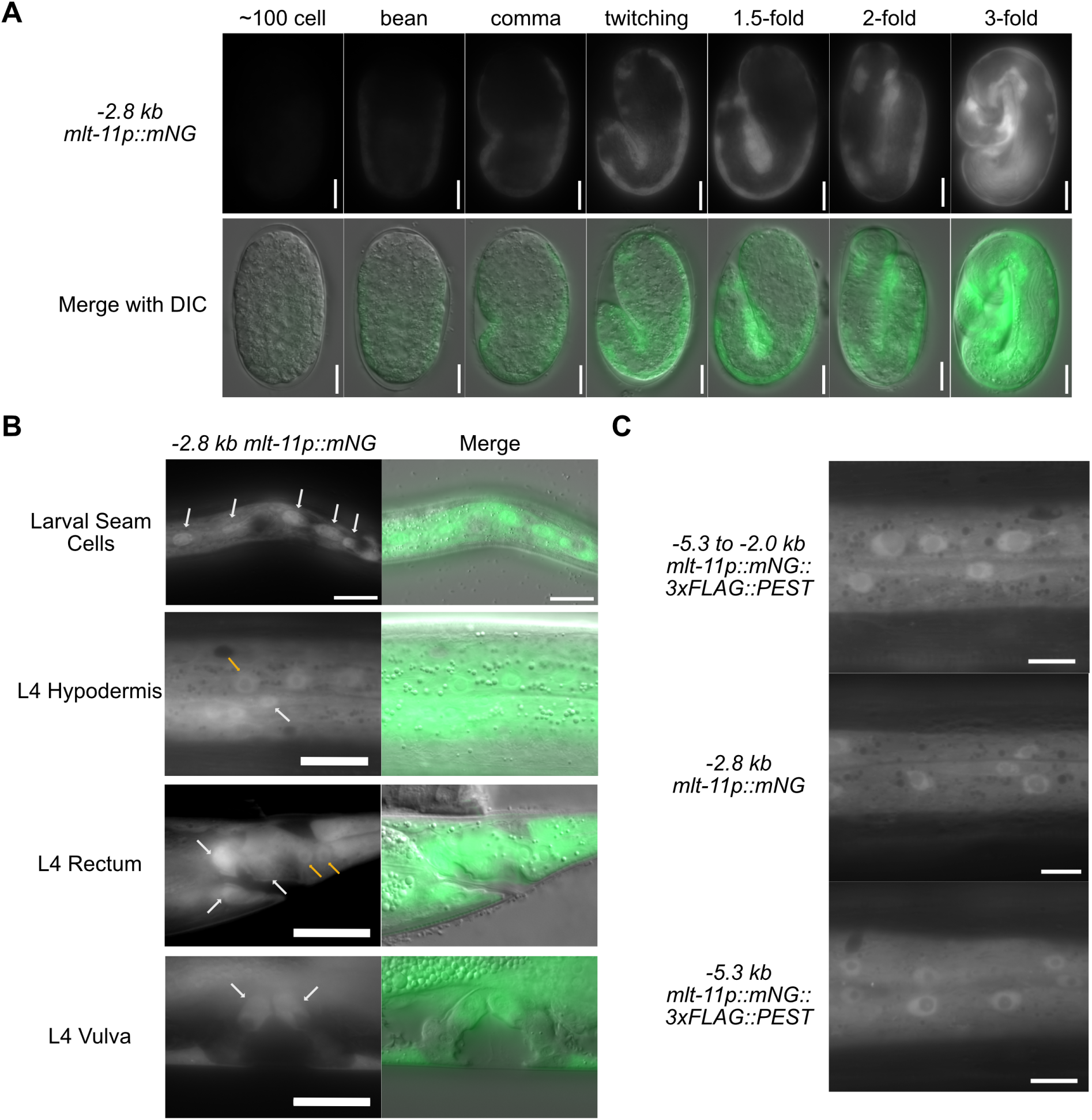
*mlt-11* is expressed in larval and embryonic epidermal cells. (A) Time course of *-2.8kb mlt-11p::mNG::tbb-2 3’UTR* expression in embryos with stages determined by embryo morphology. A minimum of 20 embryos were observed for each developmental point over two experiments. (B) Expression pattern of a *-2.8kb mlt-11p::mNG::tbb-2 3’UTR* promoter reporter in hypodermal cells of L4 worms. White arrows indicate seam cells (Larval Seam Cells and L4 Hypodermis), rectal epithelial cells (L4 Rectum) and vulval cells (L4 Vulva) and yellow arrows indicate hypodermal cells in external body cuticle (L4 Hypodermis) and near the rectum (L4 Rectum). Images are representative of 40 animals examined over two biological replicates. (C) Hypodermal expression pattern in L4s of three single-copy integrated promoter reporters. The -5.3 to -2.0 kb reporter uses the same sequence as that in Frand *et al*. (2005) and the other two use 2.8 kb and 5.2 kb of sequence upstream of the *mlt-11* transcriptional start site (TSS). Images are representative of 40 animals examined over two biological replicates. Scale bars are 10 µm for embryos in A and 20 µm in B-C.

### NHR-23 regulates *mlt-11* transcriptionally by associating with multiple regions of the *mlt-11* promoter

We next set out to identify *cis-*regulatory elements in the overlapping sequence of our 2.8 kb promoter reporter and the 5.3 kb to 2 kb reporter. There are four NHR-23 ChIP-seq peaks in the *mlt-11* promoter (Gerstein et al. 2010; Johnson et al. 2023), and the sequences under these peaks are highly conserved in other nematodes (Figure 2A; see Conservation track). There are also single NHR-23 peaks in the gene body and 3’ UTR, which we did not pursue further. There is a strong NHR-23 ChIP-seq peak (Peak 3) contained in both the 5.3 kb to 2 kb promoter reporter and our 2.8 kb promoter reporter (Figure 2A). The 5.3 kb to 2 kb reporter also contained NHR-23 ChIP-seq peak 4 (Peak 4) and our reporter also contained NHR-23 ChIP-seq peaks 1 and 2 (Figure 2A). We therefore tested whether the sequence contained in each NHR-23 ChIP-seq peak, as well as two candidate enhancers identified by ATAC-seq were sufficient to drive the expression of a reporter containing a *pes-10* minimal promoter. The *pes-10Δ::mNeonGreen* transgene displayed undetectable expression in the absence of an added *cis-*regulatory element (Figure 2B and C). When the sequences from peaks 3 and 4, respectively, were added we detected reporter expression in hypodermal cells with the peak 3 reporter having the most robust expression (Figure 2B, C). Additionally, reporters that contain the sequences from peaks 3 and 4 expressed in seam cells (Figure 2B and C). We could not detect expression of reporters containing the sequences from peak 1 nor from two ATAC-seq peaks (Figure 2B-C). The peak 2 reporter did display very low expression in seam cells though this was close to background levels (Figure 2B-C). To test whether NHR-23 regulated expression of the *mlt-11 peak 3 pes-10::mNeonGreen* promoter reporter, we performed *nhr-23* RNAi, which reduced expression of the full length *mlt-11p (-5.2kb)::mNeonGreen:: 3xFLAG::PEST* promoter reporter (Figure 2D and E) as well as *pes-10Δ::mNeonGreen* reporters containing *mlt-11* peak 3 and peak 4 (Figure 2D and E). These data indicate that NHR-23 regulates *mlt-11* and that the DNA sequences in NHR-23 ChIP-seq peaks 3 and 4 play an important role in this regulation.

**Figure 2.**
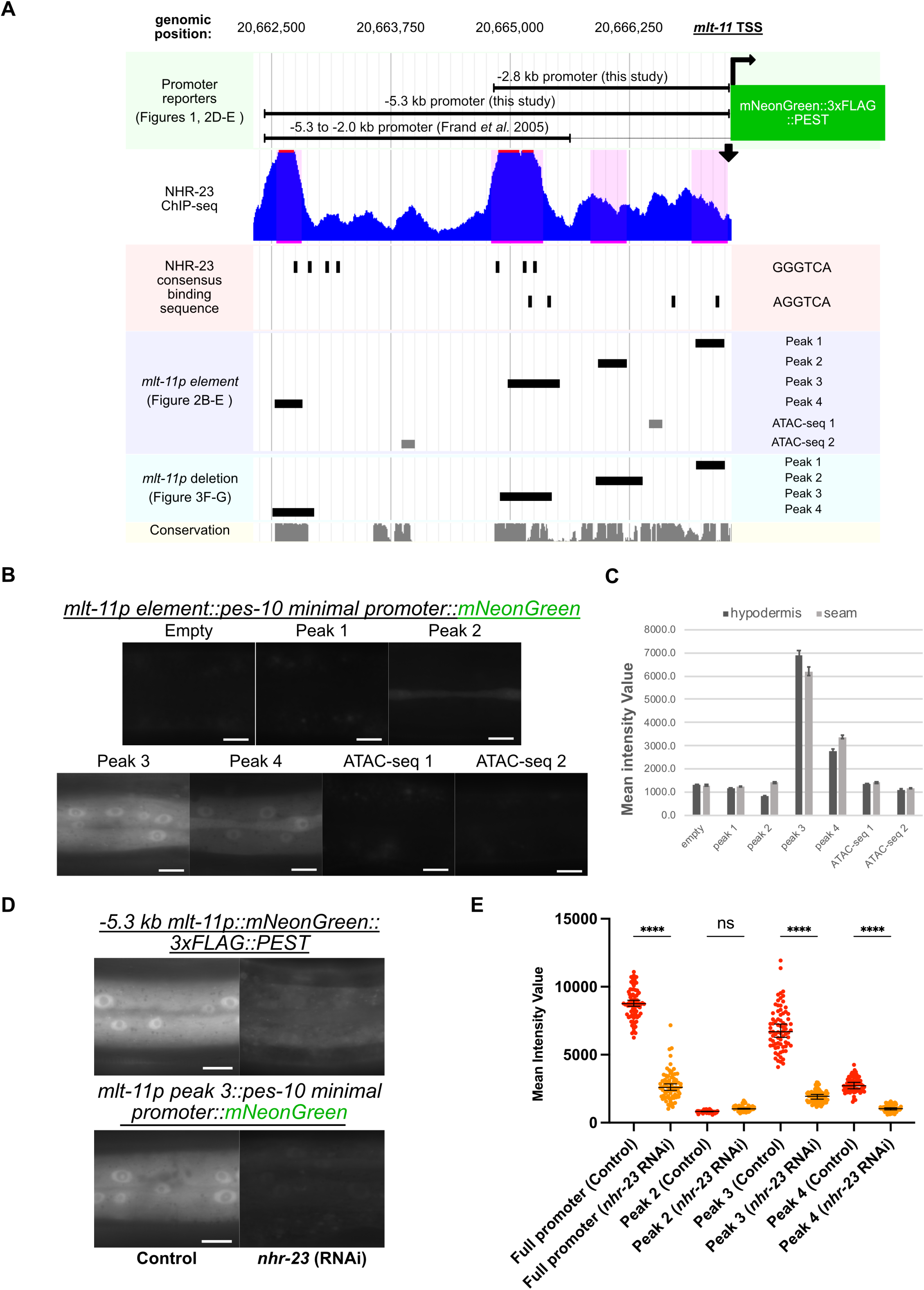
NHR-23 occupied regions in the *mlt-11* promoter are sufficient to drive reporter expression. (A) Genome browser track of relative genomic locations of *mlt-11* promoter reporters used in this study; *mlt-11* promoter depicting NHR-23 ChIP-seq peaks; relative genomic locations of NHR-23 binding motifs; relative genomic locations of candidate *cis-*regulatory elements from the *mlt-11* promoter inserted into a *pes-10* minimal promoter in this study; relative genomic locations of deletions made and tested in the *mlt-11* promoter in this study and conservation calculated across 26 nematode species. (B) Hypodermal expression pattern in L4s of single-copy integrated promoter reporters with the *pes-10* minimal promoter elements combined with *mlt-11* promoter sequence fragments corresponding to either NHR-23 ChIP-seq peaks or ATAC-seq open chromatin regions. (C) Quantification of intensity measured in hypodermal or seam cells for each promoter reporter in (B). 100 µm^2^ boxes were drawn over hypodermal cells using the Zen image processing program and intensity of pixels in the boxes measured. Statistical significance determined by one way ANOVA using Graphpad Prism 10. A minimum of 20 worms were measured for each condition over two independent experiments. (D) Hypodermal expression pattern in L4 larvae of the indicated single-copy integrated promoter reporters. Reporter strains were fed either control or *nhr-23* RNAi bacteria. (E) Quantification of intensity measured in hypodermal cells for each promoter reporter included and not shown in (D). 100 µm^2^ boxes were drawn over hypodermal cells using the Zen image processing program and intensity of pixels in the boxes measured. Statistical significance determined by one way ANOVA using Graphpad Prism 10. A minimum of 20 worms were measured for each condition over two independent experiments. Scale bars are 20 µm in B and D.

### DNA sequences in NHR-23 ChIP-seq peaks 3 and 4 are necessary for endogenous MLT-11 expression

We next wanted to test whether the NHR-23 occupied regions from the ChIP-seq dataset were necessary for endogenous MLT-11 expression with an eye to creating mutants with a range of MLT-11 expression. To monitor MLT-11 levels, we knocked an *mNeonGreen::3xFLAG* cassette into the 3’ end of the 7th *mlt-11* exon producing an internal translational fusion that labels all described *mlt-11* isoforms (Figure 3A). The strain did not display any *mlt-11* inactivation phenotypes, suggesting that the knock-in did not disrupt MLT-11 function. MLT-11::mNeonGreen::3xFLAG (MLT-11::mNG(int)) was observed in the aECM of hyp7 and seam cells at the intermolt. During the L4 larval stage, in cuticle above seam cells, localization begins at L4.3 and disappears by larval substage L4.6. In hyp7, MLT-11::mNG(int) localized first at L4.3 in thin lines reminiscent of furrows which then transitioned to two parallel lines in L4.4 separated by a gap and finally in annuli in L4.5 disappearing prior to the subsequent molt (Figure 3B). MLT-11::mNG(int) was detected consistently throughout larval development in lysosomes of the hypodermis as determined by vesicle morphology/size (Miao et al. 2020) and colocalization with a lysosomal NUC-1::mCherry fusion protein (Figure 3C; Clancy et al. 2023). MLT-11::mNG(int) was also observed in the epithelial aECM of other external-facing orifices: the buccal cavity, vulva, excretory duct and rectum (Figure 3D and E).

**Figure 3.**
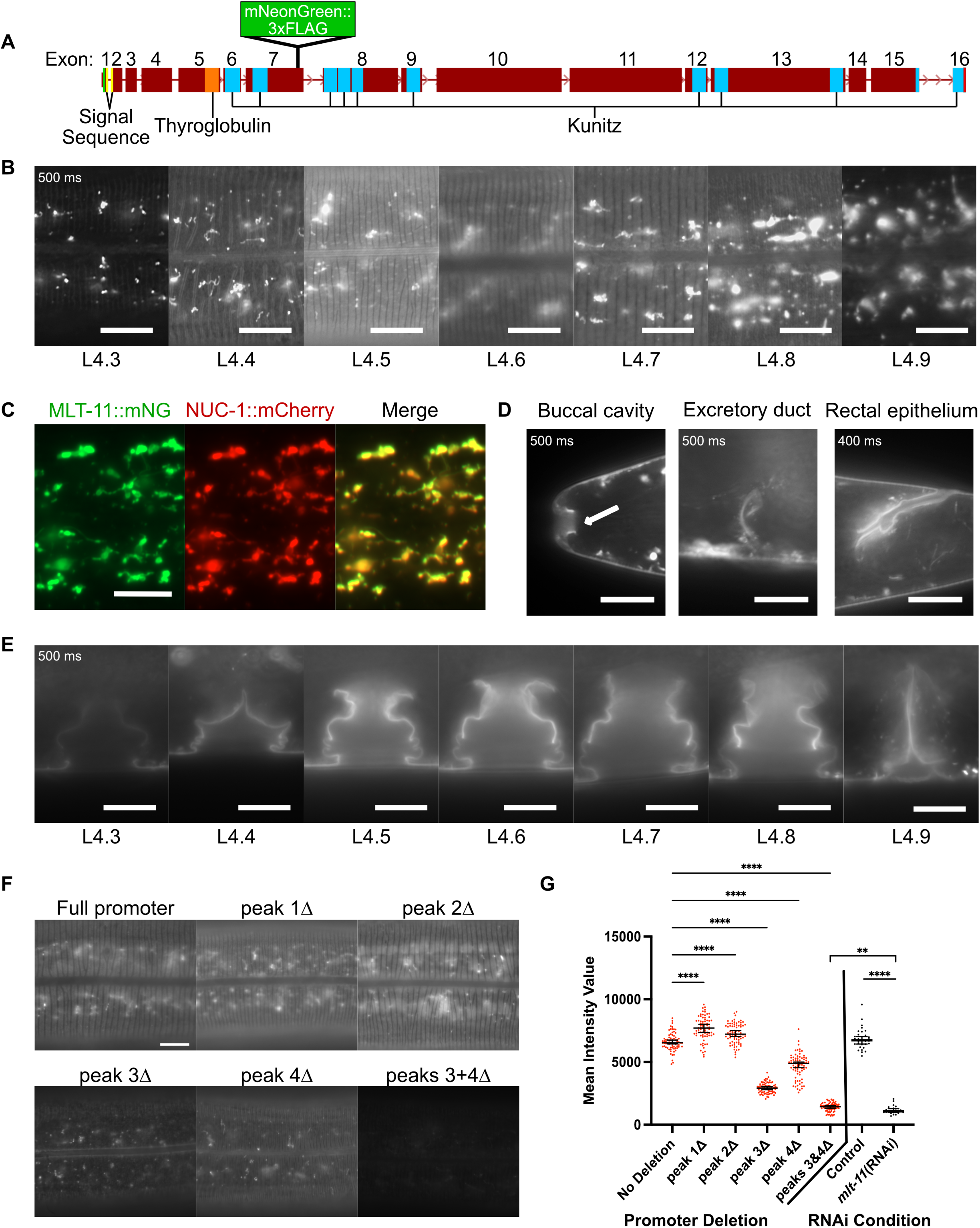
MLT-11 localizes to the aECM in seam and hypodermal cells and interfacial orifices. (A) Schematic of *mlt-11* genetic locus introns and exons, important protein domains and the internal *mlt-11::mNG::3xFLAG* knock-in. (B) Representative images of MLT-11::mNG(int) localization in hypodermal and seam cells through L4 substages. (C) Colocalization of MLT-11::mNG(int) with a NUC-1::mCherry fusion protein in lysosomes of late L4 stage larvae. (D) Representative images of MLT-11::mNG localization in the lumen of the buccal cavity, excretory duct and rectum. Images are representative of a minimum of 20 animals. (E) Representative images of vulval aECM localization of MLT-11::mNG through L4 substages. Images are representative of a minimum of 20 animals over two independent experiments. (F) Representative images of MLT-11::mNG in mid-stage L4 larvae in control animals (Full Promoter) and with endogenous deletion of the indicated promoter regions. (G) Quantification of expression of fusion proteins in this figure in the indicated genetic background or following the indicated RNAi feeding. 100 µm^2^ boxes were drawn over hypodermal cells using the Zen image processing program and intensity of pixels in the boxes measured. Statistical significance determined by one way ANOVA using Graphpad Prism 10. A minimum of 75 (promoter deletion) or 30 (RNAi Condition) measurements were made for each condition over two independent experiments. Scale bars are 20 µm in B-C and 10 µm in D-F.

Using CRISPR/Cas9, we individually deleted the sequences from each NHR-23 ChIP-seq peak in the MLT-11::mNG(int) background and observed expression of the translational fusion at substage L4.5 (Figure 3F and G). We observed the expression of MLT-11::mNG(int) to be comparable to the control following deletion of the sequences from peaks 1 or 2, but reduced with deletion of the sequences from peaks 3 and 4. Simultaneous deletion of the sequences from peaks 3 and 4 in the same strain resulted in severely reduced expression of MLT-11::mNG(int), suggesting these sequences are both necessary for full activation of expression of *mlt-11* by NHR-23. Comparing the expression reduction in the *peak 3+4Δ* mutant to *mlt-11(RNAi)* animals indicated that RNAi produced a significantly greater reduction in MLT-11::mNG(int) levels (Figure 3G). We also scored developmental rate, motility, and size (Figure 4). Interestingly, *peak 2Δ*, *peak 3+4Δ* and *mlt-11(RNAi)* animals exhibited comparable developmental delay (Figure 4A), but *peak 2Δ* animals had wild-type motility and size (Figure 4B and C). In contrast, age-matched peak 3+4Δ and *mlt-11(RNAi)* animals exhibited a significant motility defect with the RNAi causing a stronger phenotype and a comparable smaller body size (Figure 4B and C).

**Figure 4.**
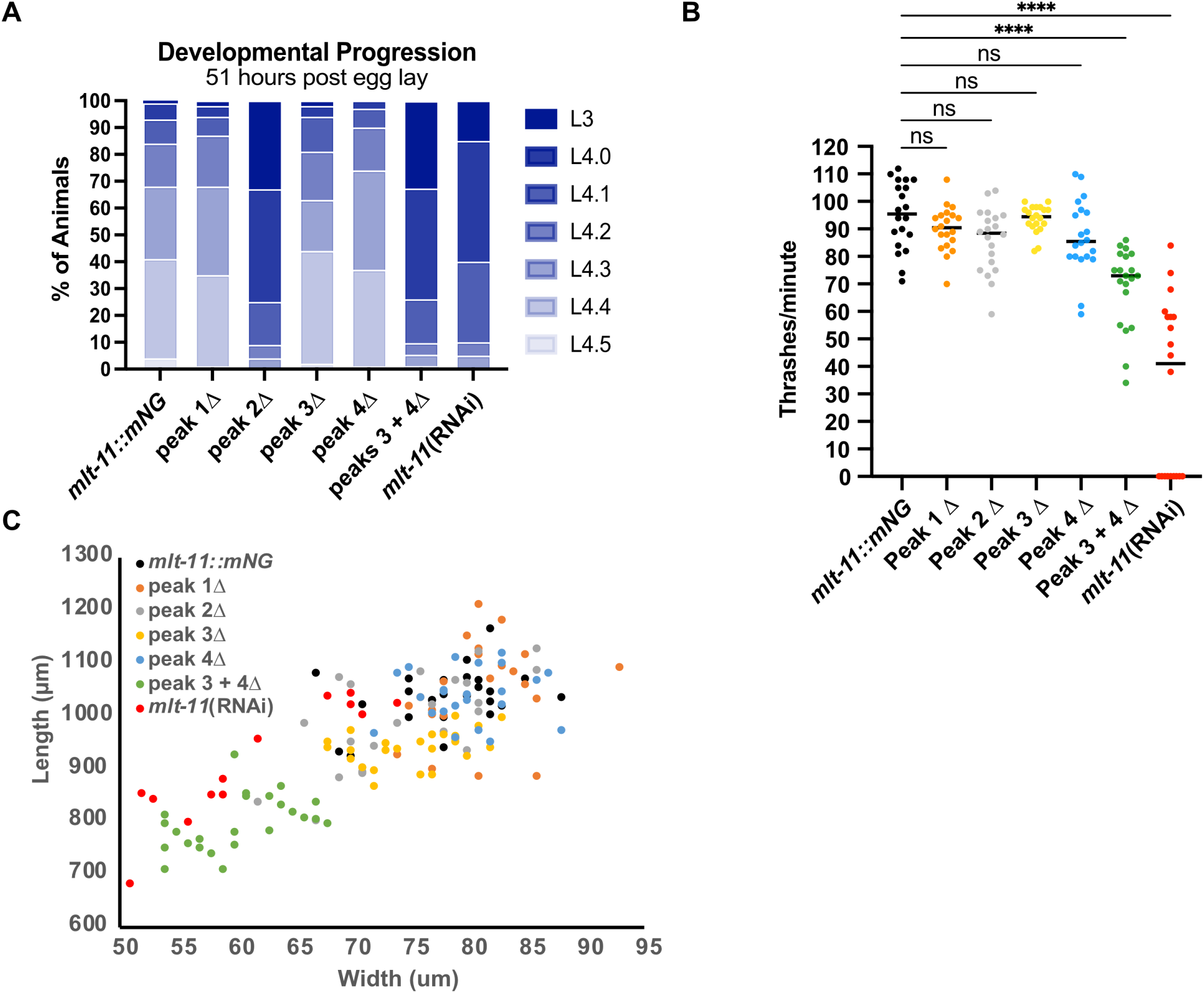
*mlt-11* expression abrogation causes developmental delay, motility defects and smaller bodies. Worms of the indicated genotypes and treatment (control or *mlt-11* RNAi) were allowed to lay eggs for two hours, then removed from the plates. Embryos were allowed to develop for 51 hours, then scored for (A) developmental stage (L3 or L4 substage), (B) thrashes per minute and (C) body length and width. Measurements represent a minimum of fifty (A) or twenty (B and C) worms over two experiments. Statistical significance determined by one way ANOVA using Graphpad Prism 10.

### Reduction of *mlt-11* expression causes defective cuticle function and structure

The developmental delay and molting defects caused by *mlt-11 (RNAi)* were reminiscent of our study of *nhr-23* (Johnson et al. 2023). As NHR-23 depletion causes a defect in the permeability barrier, we tested whether *mlt-11* inactivation also compromises this barrier. To test the cuticle barrier function, we incubated control, promoter deletion mutants and *mlt-11(RNAi)* animals with the cuticle impermeable, cell membrane permeable Hoechst 33258 dye and scored animals with stained nuclei. In wild-type control and single peak deletion animals, we observed no Hoechst staining, while we observed staining in *peak 3+4Δ* deletion mutants and *mlt-11(RNAi)* worms (Figure 5A). This barrier defect was comparable to the *bus-8* positive control strain (Partridge et al. 2008). Additionally, both *peak 3+4Δ* and *mlt-11(RNAi)* animals expressed an *nlp-29p::GFP* promoter reporter generally activated by infection, physical damage or furrow collagen inactivation (Figure 5B) (Pujol et al. 2008; Dodd et al. 2018; Martineau et al. 2021). Together, these indicate *mlt-11* is necessary for the barrier function of the cuticle.

**Figure 5.**
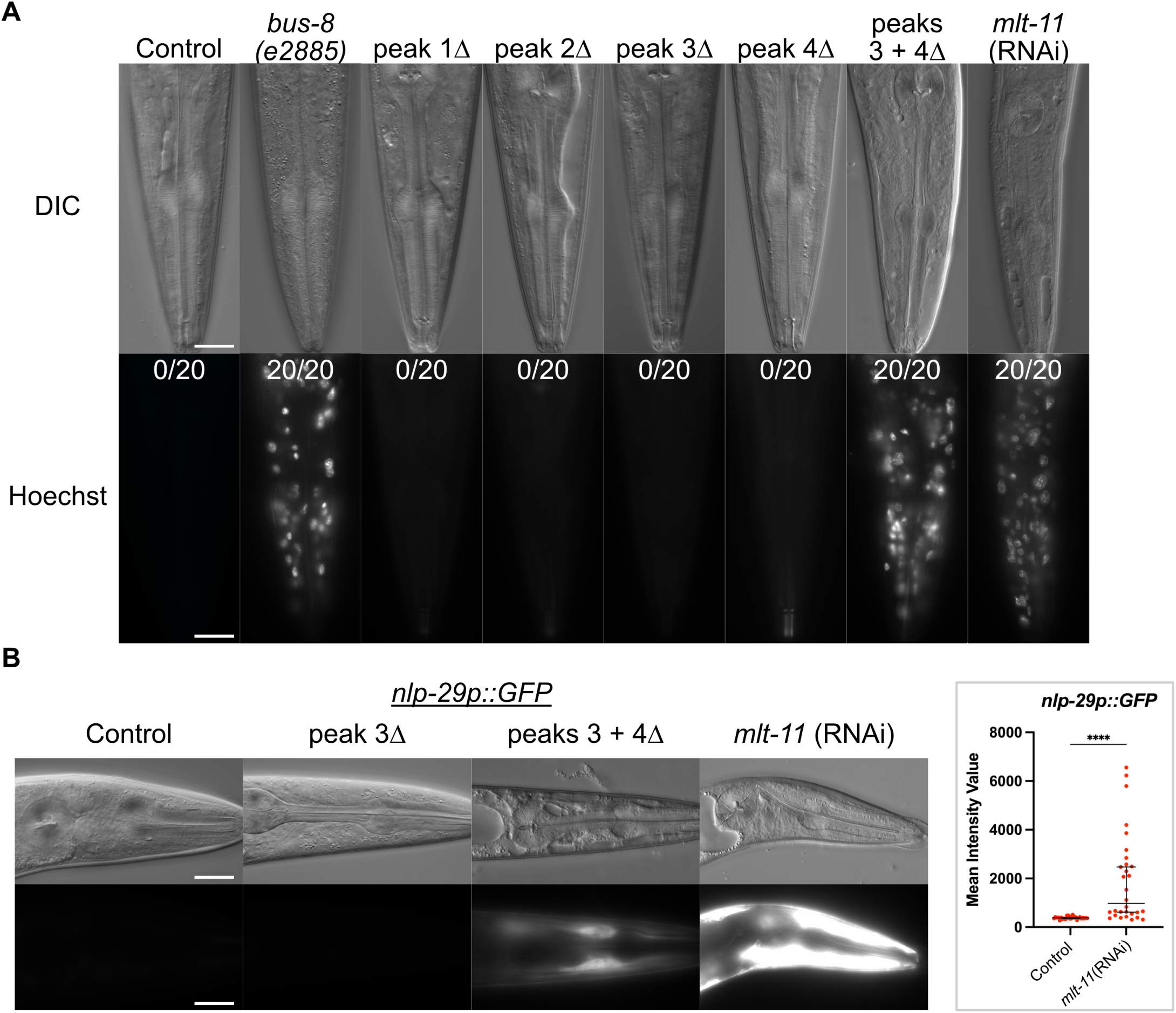
*mlt-11* knockdown causes defective aECM structure and function. (A,B) Representative images of mid-stage L4 larvae in animals of the indicated genotype/treatment. (A) Worms were washed and incubated with the cuticle impermeable/membrane permeable Hoechst 33258 dye before being imaged. A minimum of 20 worms were scored. (B) Worms carried an *nlp-29p::GFP* reporter activated by infection, acute stress, and physical damage to the cuticle (Pujol et al. 2008; Zugasti and Ewbank 2009). Three independent experiments were performed and over 40 animals scored. 100 µm^2^ boxes were drawn over hypodermal cells using the Zen image processing program and intensity of pixels in the boxes measured. Statistical significance determined by one way ANOVA using Graphpad Prism 10. Scale bars are 20 µm.

Given the barrier defect observed in *mlt-11(RNAi)* and *peak 3+4Δ* animals and that knockdown of *mlt-11* by RNAi (Frand *et al*., 2005) causes molting defects, we surmised that *mlt-11* could have a role in establishing or maintaining the structure of the cuticle. We depleted *mlt-11* by RNAi or deleted promoter elements in strains carrying translational mNG fusions to proteins that mark furrows (DPY-7, DPY-10), the basal layer (COL-19, ROL-6), medial layer (BLI-1), and cortical layer (CUT-2). We used and examined localization in late L4 animals (Figure 6A)(Peixoto and De Souza 1995; Peixoto et al. 1998; McMahon et al. 2003; Adams et al. 2023; Ragle et al. 2025).

**Figure 6.**
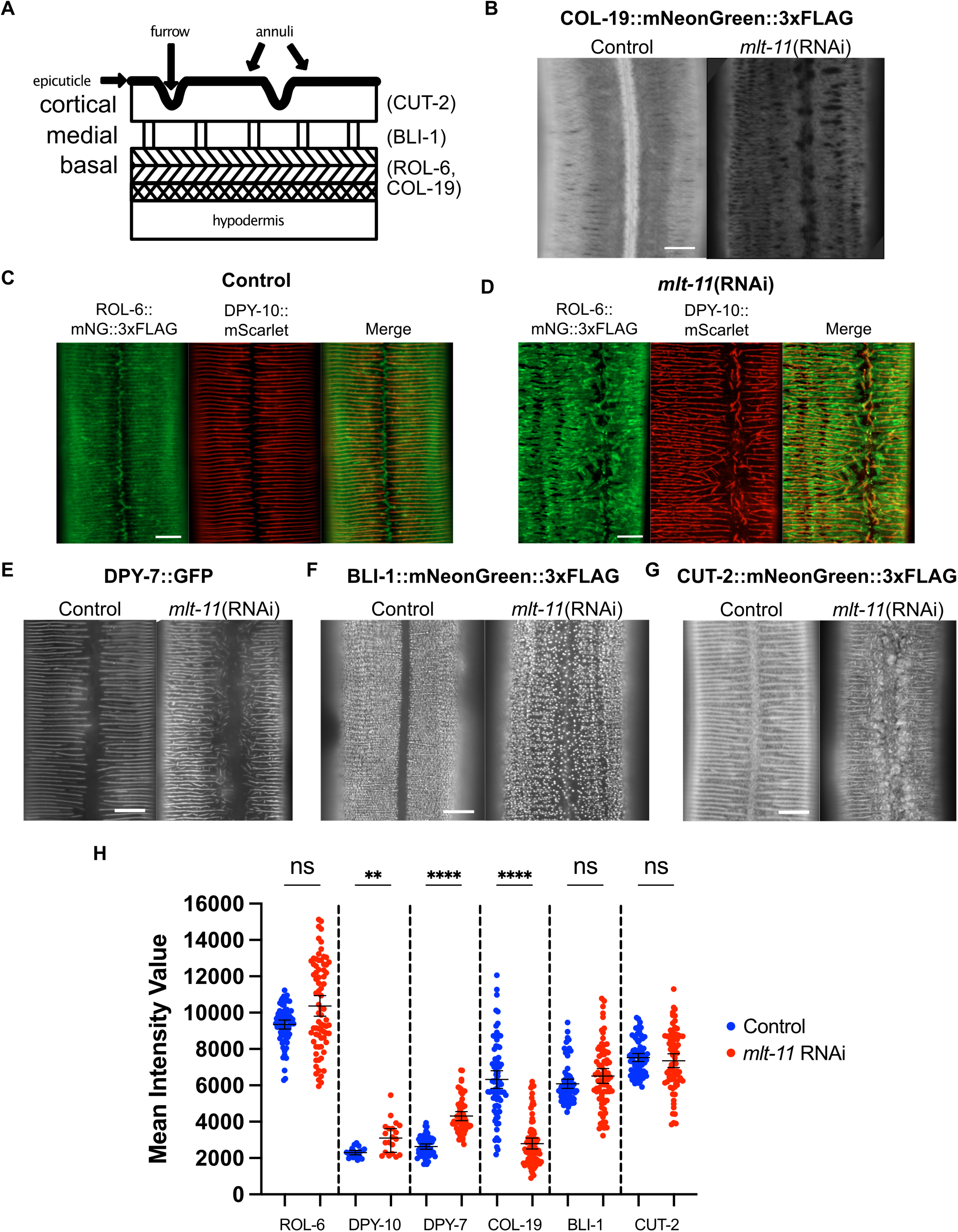
Disruption of *mlt-11* expression by RNAi or promoter deletion causes abnormalities in cuticle structure. (A) Cartoon model of the *C. elegans* adult cuticle sublayers (epicuticle, cortical, medial and basal) and topological features (annuli and furrows). The localization of some proteins used to mark specific layers of the cuticle in this study is indicated (CUT-2, BLI-1, ROL-6 and COL-19)(Ristoratore et al. 1994; Peixoto et al. 1998; Adams et al. 2023). (B) Representative images of mNG::3xFLAG fused to COL-19 in the cuticles of young adults in control or *mlt-11(RNAi)* backgrounds. (C and D) Representative images of ROL-6::mNG::3xFLAG and DPY-10::mScarlet in control or *mlt-11(RNAi)* backgrounds. (E, F and G) Representative images of DPY-7::GFP, BLI-1::mNG::3xFLAG and CUT-2::mNG::3xFLAG in the cuticles of mid-stage L4 larvae in control or *mlt-11(RNAi)* backgrounds. (H) Quantification of expression of fusion proteins in this figure. 100 µm^2^ boxes were drawn over hypodermal cells using the Zen image processing program and intensity of pixels in the boxes measured. A minimum of 20 worms were measured for each condition over two independent experiments, and the images are representative of the observed phenotype, though there was variation in severity due to RNAi penetrance phenotype. Scale bars are 10 µm in B-G.

COL-19::mNG is a basal layer marker and adult-specific collagen that normally localizes to alae and annuli, but following *mlt-11* RNAi we observed a reduction in expression and a loss of localization over the seam cells in L4 + 1 day worms (Figure 6B and H)(Thein et al. 2003). DPY-10 and ROL-6 are basal layer collagens that localize throughout L4 to longitudinal bands (furrows) and immediately adjacent to and flanking these bands, respectively (Figure 6C)(Kim et al. 2010; Johnson et al. 2023). DPY-10 and DPY-7 are members of a class of furrow collagens (DPY-2, DPY-3, DPY-7, DPY-8, DPY-9 and DPY-10); (McMahon et al. 2003; Dodd et al. 2018; Sandhu et al. 2021) that are necessary for maintaining the furrow structure of the cuticle, the correct adhesion of the cuticle to the epidermis (Aggad et al 2023) and permeability barrier function of the cuticle. ROL-6::mNG also resides in tight patches along the junction of opposing annuli above seam cells (Figure 6C). Following deletion of *mlt-11* peak 3 or peaks 3+4, ROL-6::mNG localized to both furrows and annuli through L4 (Supplementary Figure S1A). Small, but distinct gaps were observed in annuli as ROL-6::mNG localized there. Seam patches were still seen, but smaller and fewer in number. Following RNAi knockdown of *mlt-11*, annuli were punctuated with larger and more numerous gaps (Figure 6D). Localization of ROL-6::mNG was absent over seam cells and opposing annuli at the junction were completely separated from each other and often discontinuous at their termini. DPY-10::mScarlet exhibited branched and multidirectional furrow localization as well as short and discontinuous stretches of cuticle over seam cells. These stretches and the longitudinal bands were surrounded by thick clumps of ROL-6::mNG. DPY-7::mNG localized to furrows similar to DPY-10::mNG in a wild-type background and exhibited a similar branching pattern in mid-to-late L4s following *mlt-11* knockdown, suggesting *mlt-11* inactivation broadly disrupts furrow collagen localization (Figure 6E). There was a slight but significant increase in DPY-10::mScarlet and DPY-7::GFP expression following *mlt-11* RNAi (Figure 6H). BLI-1 is a structural collagen found in the medial layer of the adult cuticle, providing connection between the basal and cortical layers in the form of vertical struts (Tong et al. 2009; Adams et al. 2023). BLI-1::mNG localized in punctae organized in circumferential rows across the hyp7 cuticle, but is absent over seam cells in late stage L4s (Figure 6F). Deletion of *mlt-11* peak 3 led to gaps between the rows and inconsistency in the size of punctae (Supplementary Figure S1B). Deletion of peaks 3 and 4 together or RNAi knockdown caused further disorganization of BLI-1::mNG rows and mislocalization of punctae in the cuticle above seam cells (Supplementary Figure S1B and Figure 6F). Similarly, CUT-2, a cuticlin in the cortical layer (Lassandro et al. 1994; Ristoratore et al. 1994), had localization patterns that were altered following knockdown of *mlt-11*. CUT-2::mNG displayed aberrant localization over annuli and a fibrous pattern over seam cells in late L4s following *mlt-11* RNAi (Figure 6G). Together, this suggests *mlt-11* is necessary for proper formation or patterning of multiple layers of the adult cuticle.

Given the aberrant localization of COL-19, DPY-10, ROL-6, DPY-7, BLI-1, and CUT-2 over the seam cells, we next examined alae morphology. The lateral alae are cuticle ridges formed by the interaction between the actin cytoskeleton in epithelial cells and an extracellular provisional matrix (Cox et al. 1981; Katz et al. 2022). Three continuous longitudinal ridges span the midline of lateral surfaces of adult worms (Supplementary Figure S1C). *mlt-11(peak 3+4Δ)* animals had discontinuous alae and *mlt-11* RNAi produced a more severe phenotype. Together, these data indicate that *mlt-11* is necessary for accurate development of cuticle structures derived from both hypodermal and seam cells.

### Conserved sequences in MLT-11 play distinct roles in cuticle development

MLT-11 is predicted to be a large protein (234-341 kDA) with a signal sequence, a thyroglobulin domain, 3 lustrin domains, and 10 Kunitz protease inhibitor domains (Figure 7). A key feature of Kunitz domains is the presence of 6 conserved cysteine residues which form three disulfide bonds critical for stabilizing the domain (Ranasinghe and McManus 2013). Nine of the MLT-11 Kunitz domains contain six cysteines, while Kunitz domain 8 is missing cysteines in the second and fourth position similar to Conkunitzin-S1, a functional neurotoxin in the venom of the cone snail *Conus striatus* (Supplementary Figure S2) (Bayrhuber et al. 2005). To gain insight into MLT-11 structure and function, we generated a deletion series to determine which conserved sequences (Figure S3) were necessary for MLT-11 function. The thyroglobulin domain, Kunitz domains 9-10, a predicted furin cleavage site, and all 3 Lustrin domains appeared dispensable for development as deletion animals were viable with no obvious phenotypes (Figure 7A). Homozygous deletion of the signal sequence, the entire *mlt-11* genetic locus, Kunitz domains 2-10, 3-10, or 7-10 caused embryonic lethality (Figure 7A and Supplementary Figure S4). There was no evidence of haploinsufficiency as we could maintain balanced deletion strains. Additionally, these balanced worms produced roughly 25% arrested embryos, a rate expected for a homozygous lethal mutation (Figure S4). These data implicate the signal sequence and Kunitz domains 7-10 as being essential for embryonic development.

**Figure 7.**
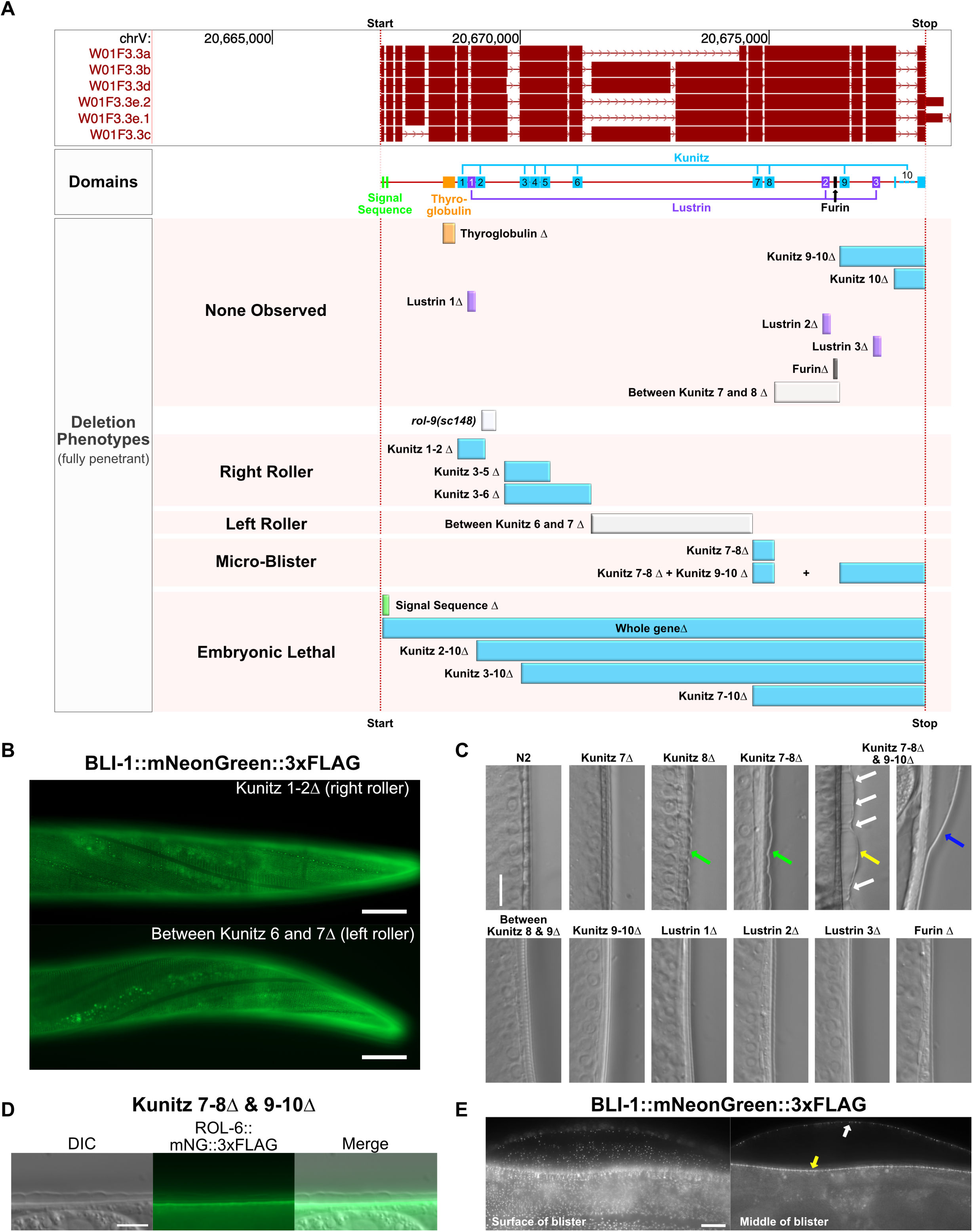
Genetically similar Kunitz domains in MLT-11 contribute uniquely to cuticle development. (A) Schematic of the UCSC Genome Browser track showing the isoforms, domains, deletions and associated phenotypes in the *mlt-11* genomic locus. (B) Representative images of BLI-1::mNG::3xFLAG expression and localization in a Kunitz 1-2Δ right roller and a strain harboring a deletion between Kunitz 6 and 7 left roller. (C) Representative images of various deletion strains showing fully intact cuticles (most strains), small µBli (green arrows), medium µBli (yellow arrows), cortical layer connections maintained with the basal layer through medial struts (white arrows), and larger blister (blue arrows). (D) Representative images of DIC and basal layer ROL-6::mNG::3xFLAG in Kunitz 7-8Δ + Kunitz 9-10Δ double deletion background. (E) Representative images of BLI-1::mNG::3xFLAG foci in the surface of a larger blister, the basal layer adjacent to the hypodermis and the cortical layer that has separated from the basal layer. (B-E) Images represent a minimum of 40 worms from a minimum of three independent experiments. Scale bars are 20 µm in B and 10 µm in C-E.

Animals with deletions spanning Kunitz domains 1-2, 3-5 and 3-6 were completely viable, producing no dead eggs as homozygotes, but instead had readily obvious (K1-2Δ) or faintly evident (K3-5Δ and K3-6Δ) right roller phenotypes (Figure 7A and B). These phenotypes are consistent with the in-frame deletion in *rol-9(sc148)* that removes one of the conserved Kunitz 2 cysteines as well as downstream sequence (Rich et al. 2022). The K1-2Δ roller phenotype was stronger than that of rol*-9(sc148).* Like *rol-9(sc148)*, the K1-2Δ roller phenotype was detectable in heterozygotes indicating that they are both dominant alleles, though the severity of the roller phenotype was more pronounced in homozygotes. In the milder roller alleles (K3-5Δ and K3-6Δ) we could not detect rolling heterozygotes. Surprisingly, deletion of the large genetic region between Kunitz domains 6 and 7 caused a left roller phenotype (Figure 7A-B).

The deletion of Kunitz 8 or the region spanning Kunitz domains 7-8 caused very small and regular separations of the cortical layer from the basal layer we have termed microblisters (µBli, Figure 7C green arrows). We observed these subtle separations across the entire cuticle from nose tip to tail. These µBli’s are larger and more frequent in worms with combined deletion of the regions spanning Kunitz 7-8 and Kunitz 9-10 (Figure 7C yellow arrow), suggesting these domains contribute to this facet of cuticle integrity. Connections between the cortical and basal layers were still maintained in these animals (Figure 7C white arrows) but lost in many cases where much larger blisters formed (Figure 7C blue arrow). To ensure these cuticle deformities were separations of distinct layers within cuticle and not separations of the entire cuticle from the hypodermis as observed in furrow collagen mutants (Aggad et al. 2023), we expressed a ROL-6::mNG fusion in the Kunitz 7-8/Kunitz 9-10 double deletion background (Figure 7D). The fusion protein localized to the basal layer immediately adjacent to the hypodermis but was absent from the cortical layer suggesting this connection was intact and these cuticle disruptions were in fact blisters. To elucidate the fate of medial layer connections in these µBli worms, we expressed a BLI-1::mNG fusion in the Kunitz 7-8/Kunitz 9-10 double deletion strain (Figure 7E). This led to an enhancement of the phenotype with blisters becoming larger and more penetrant because the *bli-1::mNG* is likely a cryptic reduction-of-function allele. BLI-1::mNG foci were seen embedded in both the cortical (Figure 7E white arrow) and basal (Figure 7E yellow arrow) sub-layers of the separated cuticle. The unique phenotypes resulting from deletion of distinct Kunitz domains suggest specific roles for these domains during cuticle development.

### MLT-11 is processed and the C-terminal fragment displays distinct localization and dynamics

Given the distinct phenotypes produced by domain deletions in the N and C-terminal portion of the protein, we returned to a C-terminal translational fusion that we had previously generated (MLT-11::mNG(C-term))(Figure 8A)(Clancy et al. 2023). Our previous analysis of this strain suggested MLT-11::mNG(C-term) was proteolytically cleaved and predominantly localized to lysosomes with weak expression in the rectum, vulva, and cuticle (Clancy et al. 2023). We also detected localization to other interfacial matrices, such as the cuticle lining the opening of the pharynx and the excretory duct (Figure 8B). Interestingly, we observed diffuse localization in the vulval lumen (Figure 8B) which sharply contrasted with the localization of the internal MLT-11::mNG to apical surfaces of vulval cells (Figure 3E). To gain insight into MLT-11 processing we performed a western blot time course (Figure 8C). We detected weak MLT-11::mNG(C-term) expression in early L4 and then strong expression of both a full-length product and a processed fragment by mid L4 (Figure 8C). Given the ∼31.5 kDa size of the mNG::3xFLAG tag, the band size is consistent with a 20-30 kDa MLT-11 C-terminal fragment produced by cleavage between Kunitz domains 8 and 9. By late L4 the bulk of product detected was cleaved MLT-11 and levels then became undetectable as MLT-11::mNG(C-term) is not expressed in adults.

**Figure 8.**
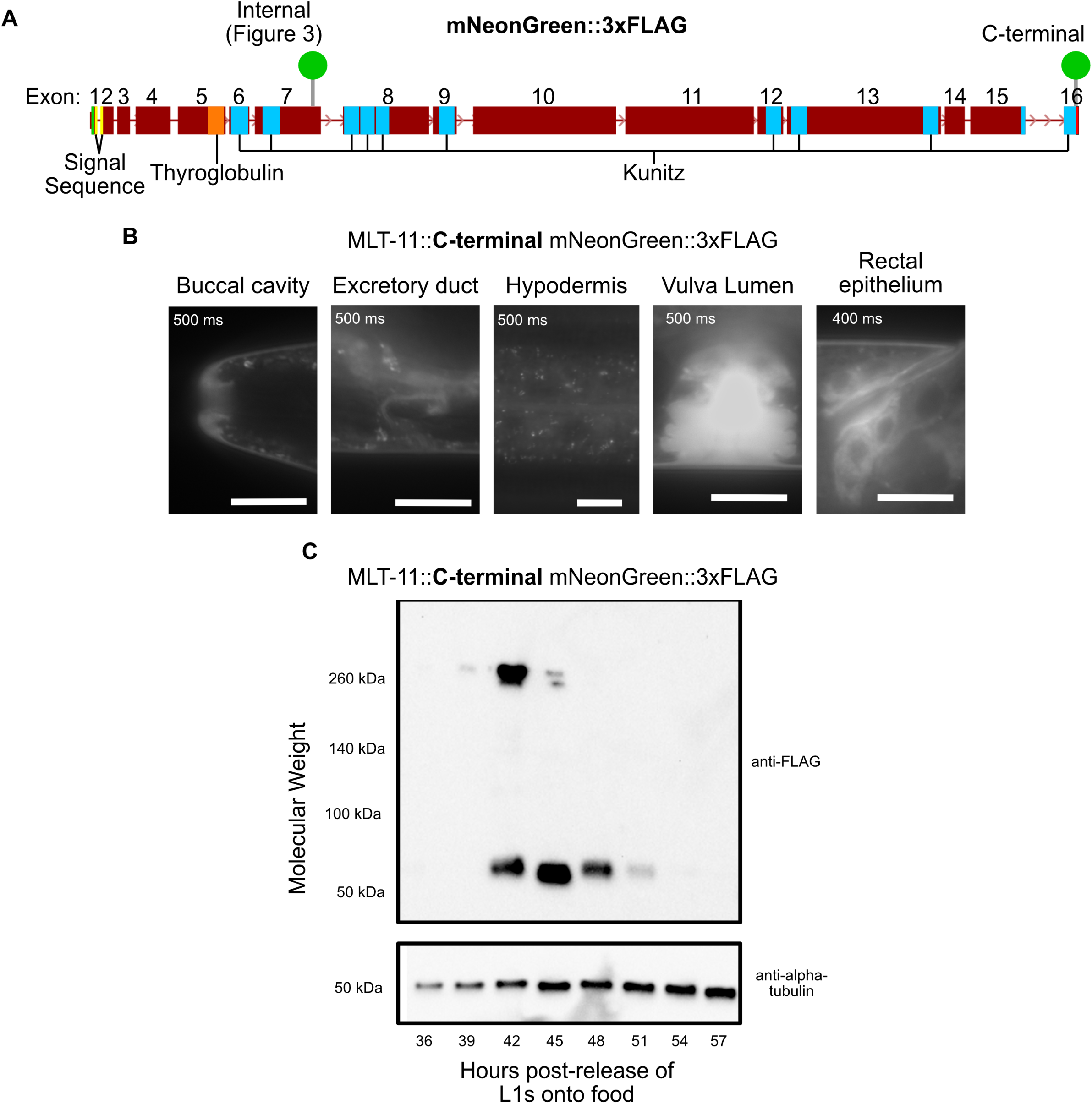
MLT-11 is processed and the C-terminal fragment displays distinct localization and dynamics. (A) Schematic of *mlt-11* genetic locus introns and exons, important protein domains and the internal and C-terminal *mlt-11::mNG::3xFLAG* knock-in. (B) Representative images of MLT-11::mNG localization in the buccal cavity, excretory duct, hypodermal/seam cells, vulva lumen and rectal epithelium. Images are representative of 40 animals examined from a minimum of three independent experiments. Scale bars are 10 µm in all images except hypodermis which is 20 µm. (C) Immunoblotting with the indicated antibodies of *mlt-11::mNG::3xFLAG* lysates harvested at the indicated time points post-release. A developmental stage for each time point as determined by vulva morphology is provided (Mok et al. 2015). Blot is representative of two independent experiments.

### *mlt-11* is essential for embryogenesis

Internal and C-terminal MLT-11::mNG translational fusions display distinct vulval localization patterns, Kunitz domains 7-10 are essential for embryogenesis, and MLT-11 is processed into at least two fragments. We therefore performed a time course to examine internal and C-terminal MLT-11::mNG localization over embryonic development. Internal MLT-11::mNG localization was observed in the embryonic sheath beginning in the bean stage and covered the entire animal until shortly before hatching. At this point, localization in punctae internal to the hypodermis and reminiscent of lysosomes was observed (Figure 9A yellow arrows). C-terminal MLT-11::mNG expression localized to the embryonic sheath in the bean stage, but also in the extraembryonic space between the animal and the egg case (Figure 9B blue arrows). This localization persisted until just prior to hatching, at which point MLT-11::mNG(C-term) was detected in the gut of many embryos (Figure 9B white arrow) as well as in internal hypodermal punctae (Figure 9B yellow arrow). These data suggest N- and C-terminal regions of MLT-11 have differing expression and localization patterns.

**Figure 9.**
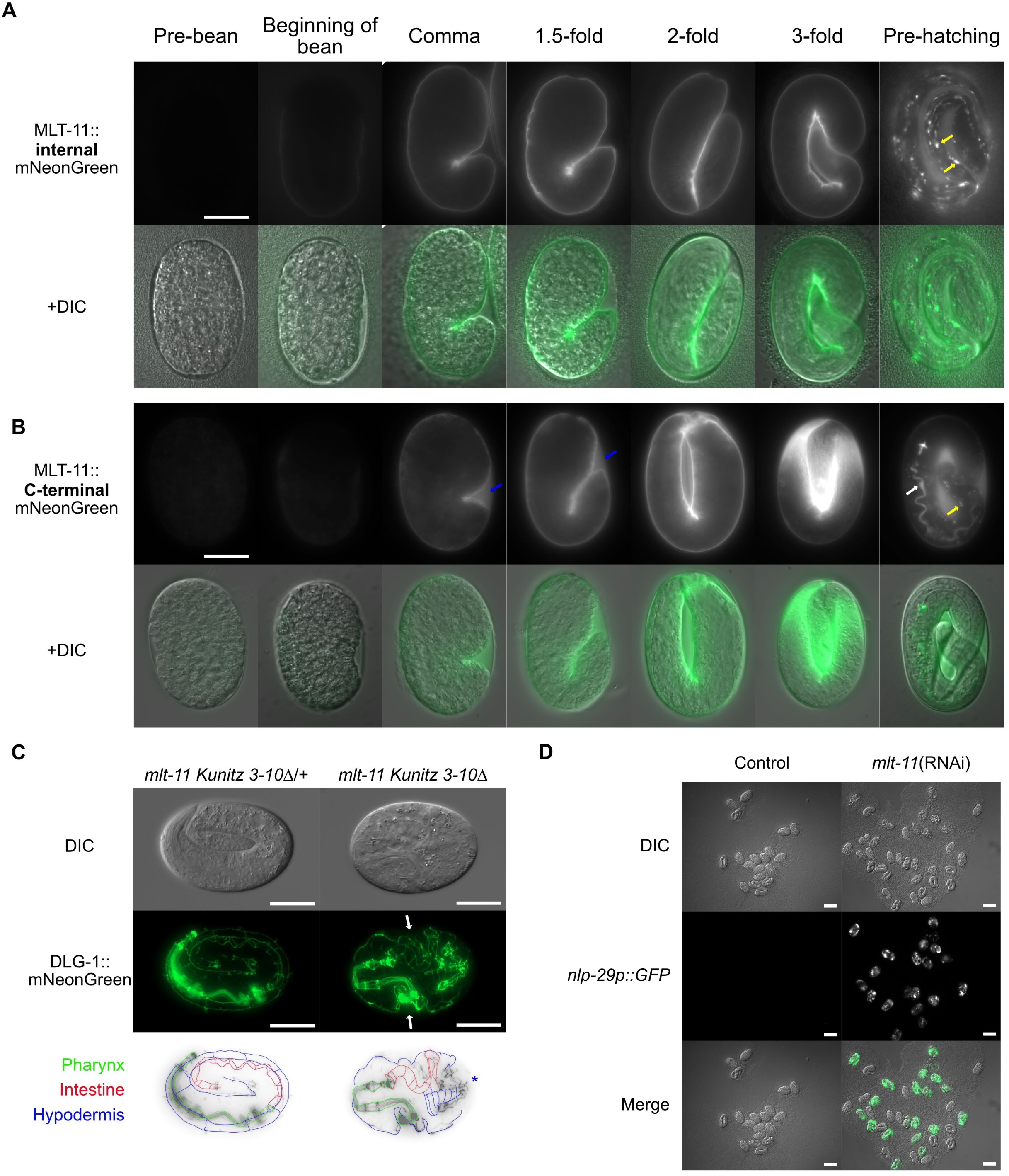
*mlt-11* is an essential gene required for embryogenesis. (A and B) Time course of internal or C-terminal MLT-11::mNG expression, respectively, in embryos with stages determined by embryo morphology. Yellow arrows indicate localization to globular structures in hypodermal cells reminiscent of lysosomes. White arrow indicates C-terminal MLT-11::mNG in the lumen of the intestine. Images represent over 20 embryos for each developmental time point in multiple observations. Blue arrows indicate C-terminal MLT-11::mNG in the extra-embryonic space between the organism and the egg case. (C) DIC and DLG-1::mNG images of embryos laid by a *mlt-11(Kunitz 3-10Δ)* heterozygous animal. One-quarter of the embryos show the disrupted DLG-1 pattern on the right. Images are representative of 40 animals examined over two biological replicates. White arrows show invaginations of the outer membrane in homozygous mutant embryos. Blue star indicates the suspected aggregation of epithelial cells in homozygous mutant embryos. Scale bars are 20 µm. (D) Images of *nlp-29p::GFP* embryos from young adult worms plated on control or *mlt-11* RNAi bacteria. Images represent two biological replicates. Scale bars are 50 µm.

The localization dynamics of MLT-11(int) were reminiscent of pre-cuticle components such as NOAH-1, which are required for maintaining embryo integrity during elongation (Vuong-Brender et al. 2017). We therefore investigated the integrity of cell membranes using a DLG-1::mNG allele to mark adherens junctions (Heppert et al. 2018). In control embryos, DLG-1::mNG labeled adherens junctions in the pharynx, intestine, and hypodermis (Figure9C). In contrast, C-terminal Kunitz deletion embryos matched to the same stage displayed severe disorganization similar to inactivation of the precuticle components *noah-1* and *noah-2* (Vuong-Brender et al. 2017)(Figure 9C). The pharynx and foregut adherens junctions appeared wild type, but the remainder of the junctions were disorganized and there was evidence of invaginations and severe disorganization in the hypodermis (Figure 9C). *mlt-11* RNAi in an *nlp-29p::GFP* promoter reporter also caused embryonic lethality, suggesting that the reporter strain might be a sensitized background. *mlt-11(RNAi*) embryos activated the *nlp-29p::GFP* reporter, suggesting damage to the developing cuticle (Figure 9D). Together these data suggest that MLT-11 is necessary for the late stages of embryogenesis.

## DISCUSSION

Despite their importance, how aECMs are built and dynamically remodeled during development and disease remains poorly understood. Using the *C. elegans* cuticle as a model aECM we show that *mlt-11* is directly regulated by the NHR-23 transcription factor. MLT-11::mNG(int) transiently localizes to the embryonic and larval aECM before endocytosis and trafficking to a compartment that is most likely lysosomal. MLT-11::mNG(int) also lines openings to the exterior (vulva, buccal cavity, excretory duct, rectal epithelium). *mlt-11* inactivation affects all three layers of the adult cuticle, indicating a broad role for MLT-11 in patterning this aECM. In agreement with these broad defects in aECM component localization, *mlt-11* inactivation causes loss of the epithelial barrier and activation of an epidermal damage reporter. MLT-11 is processed into at least two fragments and internal and C-terminal mNG knock-ins display distinct localization patterns. Alleles predicted to cause a loss of MLT-11 function cause embryonic lethality with severe disorganization of hypodermal adherens junctions. Together, this work suggests that MLT-11 acts similarly to pre-cuticle components, promoting patterning of all layers of the cuticle and maintaining epithelial integrity during embryonic elongation.

### Structure function analysis suggests Kunitz domains confer functional specificity

As *mlt-11* null alleles cause embryonic lethality (Figure 7), we explored whether deletion of putative *cis-*regulatory elements in the *mlt-11* promoter could provide viable reduction-of-function alleles. *mlt-11* is an NHR-23-regulated gene with four potential NHR-23 regulatory elements in the promoter (Frand et al. 2005; Kouns et al. 2011; Johnson et al. 2023) (Figure 2). Consistent with the reporter data (Figure 2), deletion of the endogenous peak sequences revealed that peaks 3 and 4 produced the strongest effect on MLT-11::mNG(int) expression and a double deletion of peaks 3 and 4 results in even stronger reduction in expression (Figure 3). Comparing *cis*-regulatory element deletion mutants to *mlt-11(RNAi)* animals revealed a range of phenotypic severity. ROL-6::mNG and BLI-1::mNG localization appeared to be the most sensitive to MLT-11 levels, as we saw aberrant localization for both in the Peak 3 deletion mutant. The only other phenotype that we observed for a single peak deletion mutant was the curious developmental delay of the peak 2 deletion mutant, which displayed no other defects. Interestingly, a promoter reporter for this *cis-*element was expressed in the seam cells (Figure 2), and tissue-specific RNAi experiments indicate that NHR-23 activity is particularly important in seam cells (Johnson et al. 2023). An important future direction is to explore the tissue-specificity of MLT-11 activity and the roles of the seam cells and hypodermis in molting.

Peak 3+4 deletion caused a significant reduction in MLT-11 levels (Figure 3), and resulted in comparable developmental delay, smaller body size, and cuticle barrier defects to *mlt-11* RNAi (Figure 4, 5) suggesting these phenotypes are next most sensitive to MLT-11 levels. Cuticle barrier defects, *nlp-29p::GFP* reporter activation and alae gaps were caused by both Peak 3+4 deletion and *mlt-11* RNAi, though stronger phenotypes were seen in *mlt-11(RNAi)* animals (Figure 5 and Supplementary Figure S1). Ecdysis defects, specifically entrapment in partially or fully separated cuticles, were only observed following *mlt-11* RNAi, and embryonic lethality was only observed in *mlt-11* null alleles or *mlt-11* RNAi on *nlp-29p::GFP* animals, which is likely a sensitized background. (Figure 7) indicating that very low levels of MLT-11 are required to prevent these defects.

### MLT-11 is proteolytically processed and distinct activities reside in the N and C-termini

Western blotting revealed that MLT-11 is rapidly processed into a 20-30 kDa fragment, which is consistent with cleavage between Kunitz domains 8 and 9, and C-terminal and internal translational fusions displayed distinct localization patterns and dynamics (Figure 3, Figure 8, and Figure 9). We have not been successful with western blots to assess internal MLT-11::mNG abundance and size, so it is unclear whether the protein is processed into additional fragments. The internal MLT-11::mNG localization was reminiscent of pre-cuticle components, such as NOAH-1 (Vuong-Brender et al. 2017). MLT-11::mNG(int) was secreted and endocytosed in both embryos and larvae and localized to the embryonic sheath (Figure 3, Figure 7, and Figure 9). Yet curiously, deletions of Kunitz domains in this region did not produce the lethality typical of pre-cuticle components (Kelley et al. 2015; Vuong-Brender et al. 2017; Sundaram and Pujol 2024). Rather, deletions of Kunitz 1-2, 3-5, and 3-6 produced roller phenotypes. In contrast, C-terminal deletions (Kunitz 7-10) produced embryonic lethality and loss of epithelial cell integrity reminiscent of *noah-1* mutants (Vuong-Brender et al. 2017). The C-terminal translational fusion displayed a similar localization to precuticle components in embryos (Vuong-Brender et al. 2017), and also had a diffuse localization in both embryos and the vulva (Figure 8B and Figure 9A). The foregut localization of MLT-11::mNG(C-term) at the end of embryogenesis is reminiscent of BLI-4::sfGFP localization (Birnbaum et al. 2023). BLI–4 is a subtilisin/kexin family protease required for embryonic elongation, larval development, and formation of BLI-1/BLI-2 nanoscale collagen struts (Thacker et al. 1995; Adams et al. 2023; Birnbaum et al. 2023). Exploring whether MLT-11 and BLI-4 interact and whether MLT-11 impacts the BLI-4-dependent processing of collagens such as SQT-3 and DPY-17 are a logical extension of this work.

When we deleted Kunitz domains 7-8 we observed a novel “µBli” phenotype, in which the cortical and basal layer periodically separated. Like the Bli phenotype of *bli-1* or *bli-2* mutants (Adams et al. 2023), the µBli was only observed in adults suggesting that it likely arises from destabilization of BLI-1/BLI-2 struts. Supporting this assertion, BLI-1::mNG is known to be a cryptic reduction-of-function allele (Adams et al. 2023) and deleting Kunitz 7-8 in a *bli-1::mNG* background exacerbated the µBli phenotype. Deletion of Kunitz domains 9-10 also exacerbated this phenotype, leading to larger microblisters. Curiously, deletion of Kunitz domains 7-10 causes embryonic lethality, but deletion of the intervening sequence between Kunitz domain 8 and 9 was superficially wild type. It is not currently clear whether the µBli and embryonic lethality represent common molecular defects or are distinct defects.

Loss of Kunitz domains 1-2, 3-5 or 3-6 produced right rollers. These data are consistent with the molecular identification of the semi-dominant right roller *rol-9(sc148)* allele as an in-frame deletion in *mlt-11* that removes the last cysteine of Kunitz domain 2 as well as 67 amino acids of downstream conserved sequence (Rich et al. 2022). Kunitz 1-2 deletion produced a stronger semi-dominant roller phenotype, suggesting that the *sc148* domain may disrupt Kunitz 2 function. Testing the individual contributions of Kunitz 1 and 2 and engineering mutations predicted to disrupt Kunitz activity are key future experiments. Surprisingly, deletion of the sequence between Kunitz domains 6 and 7, which contained conserved sequence blocks (Supplementary Figure S3) but no predicted domains, caused a completely penetrant left roller phenotype. *mlt-11* thus appears to be in an exclusive class of “ambidextrous” roller genes in which specific mutations can produce either left or right rollers, depending on the mutation. *sqt-1* is another member of this class. Mutations in *sqt-1* that alter a conserved carboxyl domain cysteine prevent crosslinking into dimers, tetramers, and oligomers, leading to left rollers (Yang and Kramer 1999). Mutation of a predicted subtilisin protease cleavage site causes extra sequence to be retained on the SQT-1 N-terminus and a right roller phenotype (Yang and Kramer 1999). The *mlt-11* phenotypes suggest that different Kunitz domains or conserved sequences may control interactions with specific substrates. Given the critical role of proteases such as BLI-4 and DPY-31 in collagen processing, testing for genetic interactions between *mlt-11* mutations and *bli-4* and *dpy-31* alleles is an important future direction. Additionally, testing how left and right *mlt-11* roller alleles affect the localization and processing of collagens with Rol phenotypes is another priority.

### MLT-11 is required for patterning and function of multiple cuticle layers

Mutation or depletion of oscillating collagens has been shown to affect the structural organization of collagens with similar temporal expression dynamics, suggesting these collagens might be part of a common substructure (McMahon et al. 2003). Temporally, during the L4 stage, *mlt-11* mRNA peaks in expression close to when *bli-1* mRNA is expressed and before the *rol-6* and *cut-2* mRNA expression peak (Meeuse et al. 2020) and an internally tagged MLT-11::mNG translational fusion displays aECM localization from L4.3 to L4.8 with a peak localization at L4.5 (Figure 3). Surprisingly, *mlt-11* inactivation affected the localization of proteins in the basal, medial, and cortical layer of the cuticle (Figure 6) with a range of defects. In the basal layer, MLT-11 was required for the wild-type localization of all four markers tested (COL-19, DPY-7, DPY-10, ROL-6)(Figure6B-E). For COL-19, *mlt-11* RNAi caused a significant reduction in expression as well as a loss of localization to alae (Figure 6B). *mlt-11* inactivation caused the formation of short fragments of the furrow collagens DPY-7 and DPY-10 that were often at aberrant angles deviating from the wild-type circumferential pattern (Figure 6C, D, and E). It is unclear whether the shorter fragments reflect breakage of longer fibers or formation of shorter fibers. Curiously, DPY-10 fibers normally terminate over the seam cells, but following *mlt-11* RNAi short fibers formed in parallel to the longitudinal axis (Figure 6C and D). ROL-6, which normally flanks furrow collagens, displayed a disorganized aggregated pattern but still appeared to flank DPY-10 fibers (Figure 6D and E). In the medial layer, *mlt-11* inactivation resulted in larger, disorganized BLI-1 punctae that were no longer excluded from the alae region (Figure 6F). The cortical marker, CUT-2 normally forms circumferential furrow fibers and localizes over seam cells (Figure 6E). Following *mlt-11* RNAi we still observed furrow fibers, but they were frequently disorganized and there was severe disorganization flanking and above the seam cells (Figure 6G). Interestingly, NHR-23 depletion also causes aberrant ROL-6 and NOAH-1 localization over the seam cell (Johnson et al. 2023). Exploring the relative roles of seam and hypodermal cells in aECM assembly and whether these defects reflect a role for MLT-11 in patterning each layer or whether a defect in an early aECM structure caused by *mlt-11* inactivation propagates to subsequent layers are important future questions.

*mlt-11* inactivation also produced a defective cuticle barrier and activation of an epidermal stress reporter (Figure 5). It is not clear what is leading to the cuticle barrier defect following reduction of MLT-11 levels. RNAi screens of collagens found that only knockdown of furrow collagens caused a barrier defect (Sandhu et al. 2021) and epidermal stress reporter activation (Dodd et al. 2018), and *mlt-11* inactivation caused defective localization of the furrow collagens DPY-7 and DPY-10 (Figure 6C, D, and E). *bus-8* is a glycosyltransferase required for localization of the molting regulator MLT-8 to lysosomes (Wu et al. 2022). Given the localization of MLT-11 in lysosomes, it is possible that it functions in this compartment to promote barrier integrity. Alterations in the cortical layer and epicuticle can also lead to barrier defects (Njume et al. 2022; Pooranachithra et al. 2024). The dynamic aECM localization, broad effects on cuticle structure, and role in embryonic morphogenesis are highly reminiscent of pre-cuticle components like NOAH-1 (Vuong-Brender et al. 2017; Cohen et al. 2020b). Exploring the interrelationship between MLT-11 and precuticle components will provide insight into how the aECM is assembled.

MLT-11 has 10 predicted protease inhibitor domains most of which contain residues necessary for Kunitz inhibitor activity (Ascenzi et al. 2003; Ranasinghe and McManus 2013), which could buffer reduction of levels. The position and the number of Kunitz domains is strongly conserved out to parasitic nematodes with 300-500 million years of divergence from *C. elegans.* It is possible that *mlt-11* mutant and knockdown phenotypes result from aberrant protease activity. MLT-11 localizes to both the aECM and lysosomes, so phenotypes could arise from aberrant protease activity in either compartment. Some proteases, such as BLI-4 (PCSK family), SURO-1 (zinc carboxypeptidase), and DPY-31 (astacin metalloprotease) are thought to be involved in collagen processing in the secretory pathway, while others such as NAS-36 and NAS-37 (astacin metalloproteases) promote apolysis (Thacker et al. 1995; Davis et al. 2004; Suzuki et al. 2004; Frand et al. 2005; Stepek et al. 2010b; Kim et al. 2011; Stepek et al. 2011; Birnbaum et al. 2023; Sung et al. 2025). Interestingly, SURO-1 was discovered in a genetic screen that identified suppressors of a dominant left roller phenotype caused by overexpression of *rol-6(su1006)* (Kim et al. 2011). ADM-2, the sole *C. elegans* member of the ADM-meltrin metalloprotease family suppresses the molting defects in reduction-of-function *nekl* alleles that reduce endocytosis (Joseph et al. 2022). The ADAMTS protein, ADT-2, is implicated in body size control and collagen organization, which is interesting given the smaller sized animals and cuticle disorganization produced by *mlt-11* inactivation (Figure 4 and Figure 6) (Fernando et al. 2011). Cathepsins are lysosomally localized proteases (Britton and Murray 2004; Miedel et al. 2012) and lysosome dysfunction can cause impaired degradation of endocytosed aECM components and molting defects (Miao et al. 2020). Kunitz domains typically inhibit serine metalloproteases and the above-mentioned proteases are all in other protease families. However, an atypical and functionally divergent family of Kunitz domain containing proteins in the parasitic helminth, *F. hepatica* was shown to have no activity against serine proteases but rather inhibited cysteine proteases such as cathepsins (Smith et al. 2020). Thus, it is important to consider a broad range of potential MLT-11 substrates. It is also possible that MLT-11 is acting as a scaffold or accessory factor, as a related Kunitz domain containing protein, BLI-5, was shown to enhance the activity of porcine pancreatic elastase and bovine pancreatic alpha-chymotrypsin (Stepek et al. 2010a). Screening for genetic and protein-protein interactions between *mlt-11* and proteases is a high-priority future direction to determine how MLT-11 ensures proper cuticle structure/function.

## MATERIALS AND METHODS

### Strains and culture

*C. elegans* were cultured as originally described (Brenner 1974), except worms were grown on MYOB media instead of NGM. MYOB agar was made as previously described (Church et al. 1995).

### Strains created by injection in the Ward Lab and used in this study

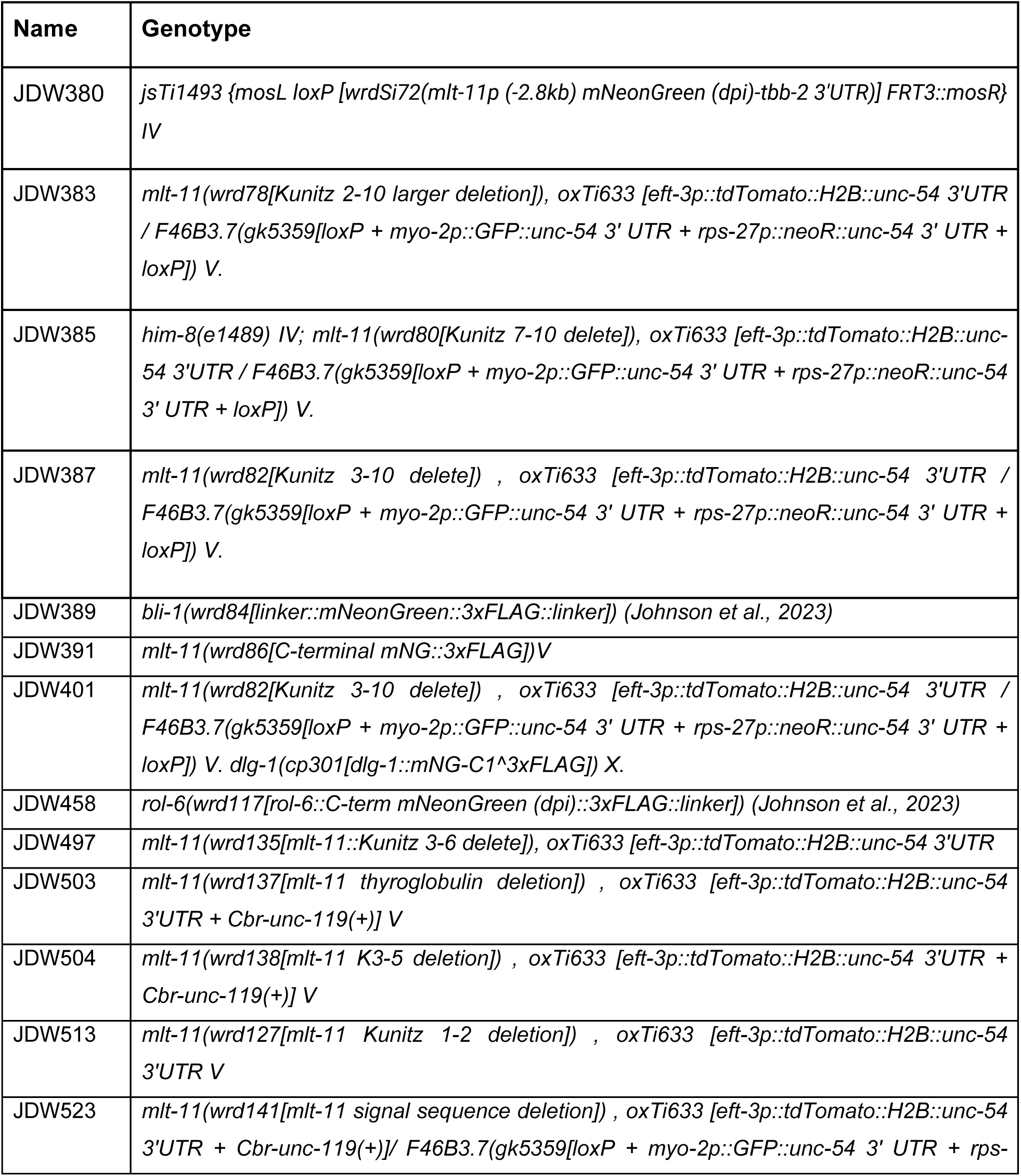
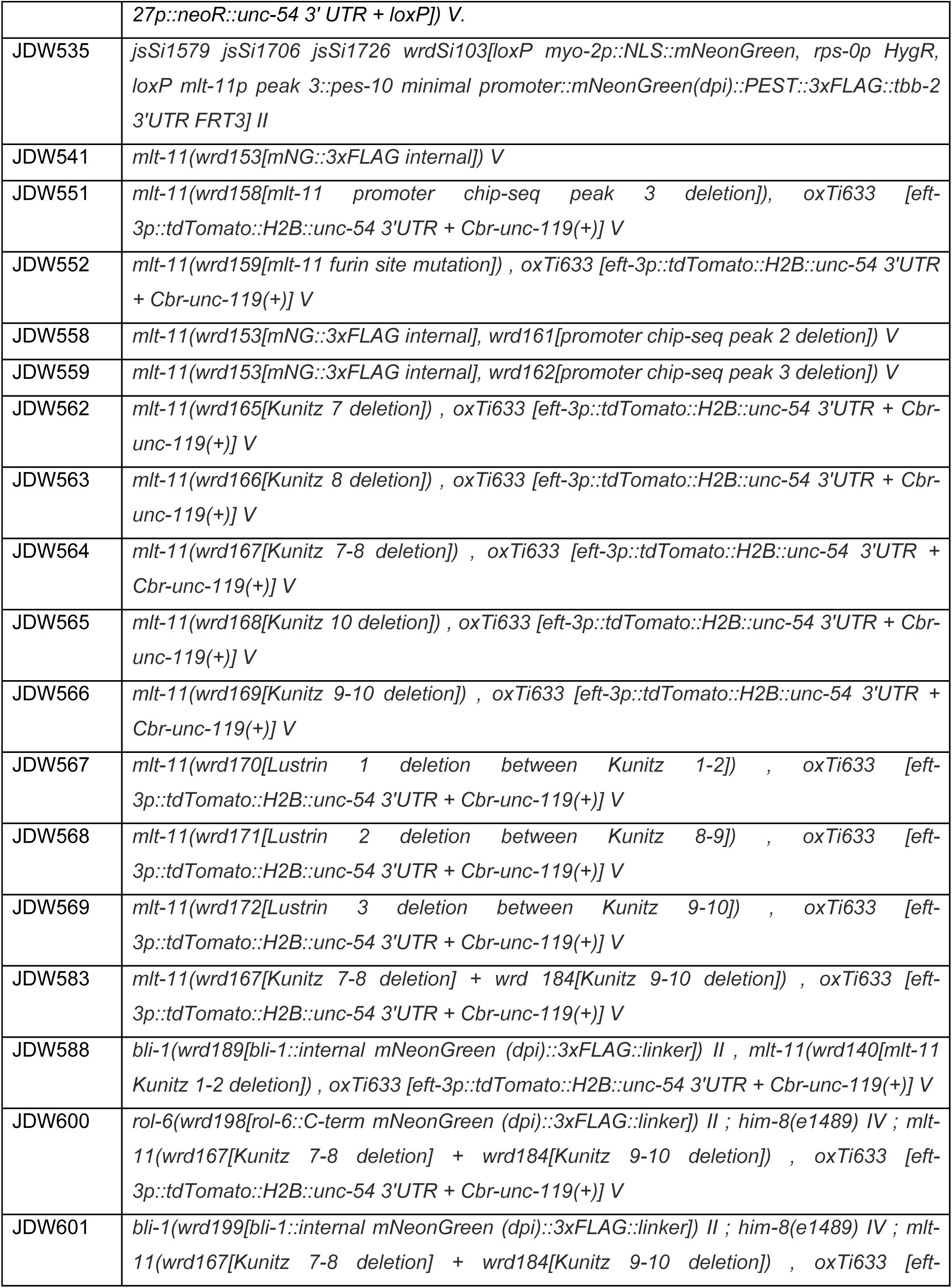
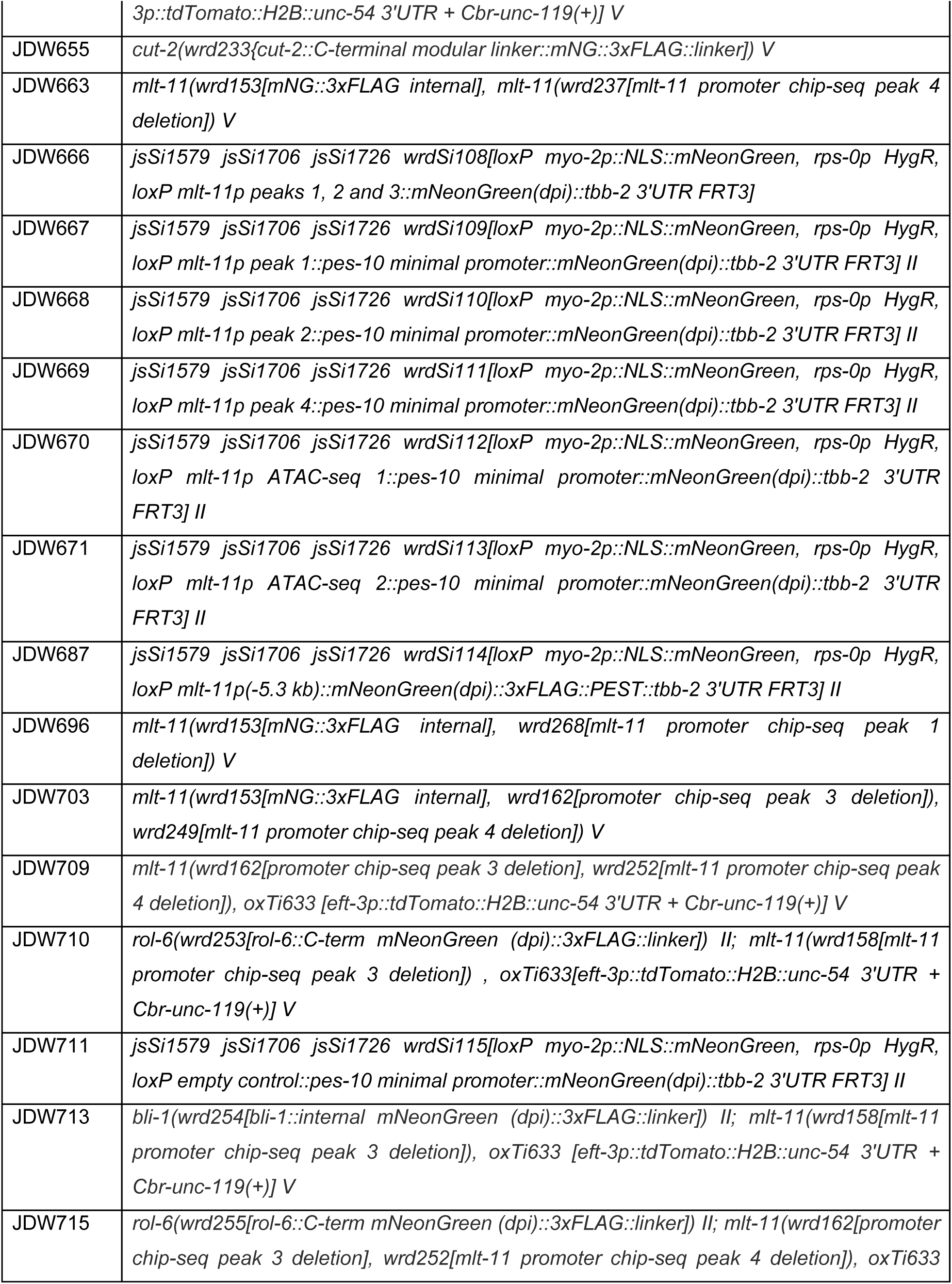
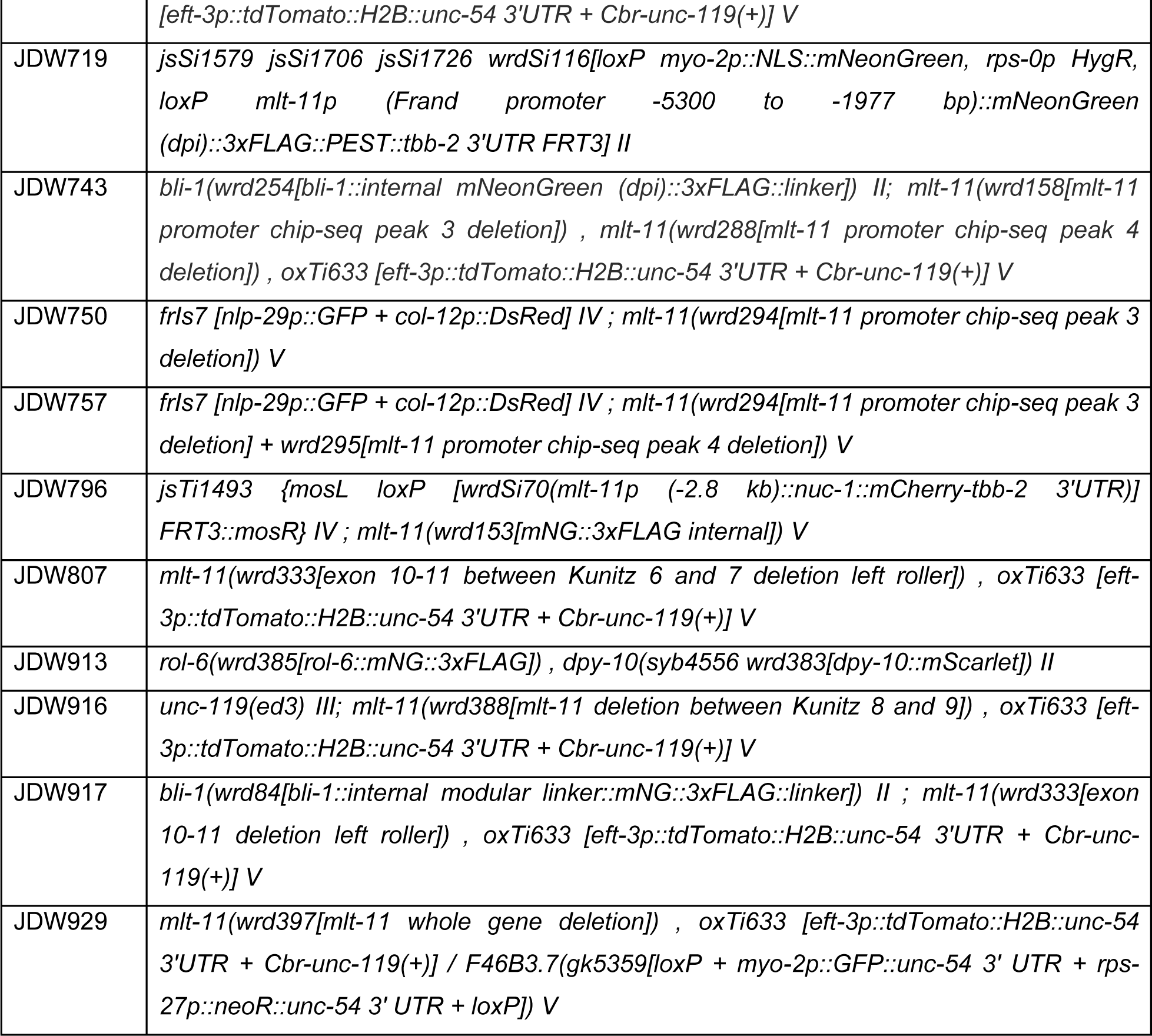

### Strains provided by the *Caenorhabditis* Genetics Center

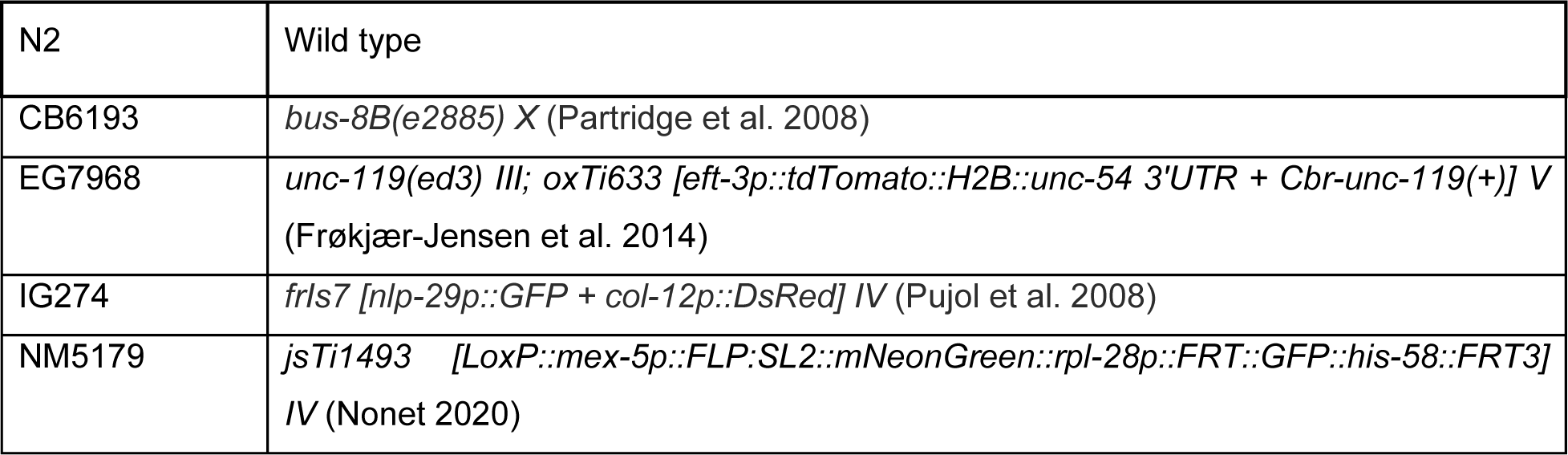
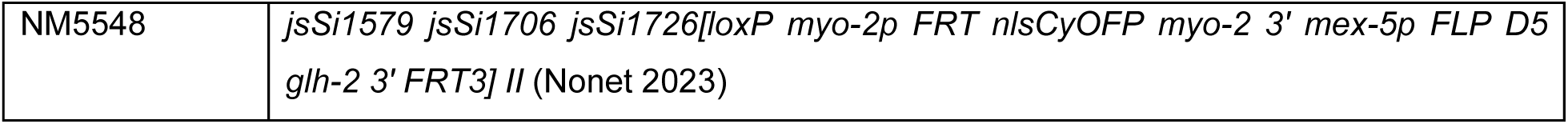

### Other strains

JDW655 *cut-2(wrd233{cut-2::mNG::3xFLAG]) V* and PHX4625 *col-19::mNG(cyb4625) X* are described elsewhere (Ragle et al. 2025). JDW913 was created by generating a *rol-6::mNG::3xFLAG* knock-in as previously described (Johnson et al. 2023) in a JDW909 *dpy-10(syb4556 wrd383[dpy-10::mScarlet]) II* background. JDW909 will be described elsewhere. XW18042 *qxIs722(dpy-7p::DPY-7::sfGFP)* was created by the lab of Xiaochen Wang (Miao *et al* 2020) and provided to us by Prof. Meera Sundaram.

### Genome Editing

All plasmids used are listed in Supplementary Table S1. Annotated plasmid sequence files are provided in File S1. Sequence files for knock-ins, promoter deletions, and coding sequence deletions are provided in File S2. Specific cloning details and primers used are available upon request. JDW380 *jsTi1493 {mosL loxP [wrdSi72(mlt-11(-2.8kb)p:: mNeonGreen(dpi)::tbb-2 3’UTR)] FRT3::mosR} IV* was created by recombination-mediated cassette exchange (RMCE)(Nonet 2020). A 2.8 kb *mlt-11* promoter fragment was initially Gibson cloned into the *NLS::mScarlet (dpi)::tbb-2 3’UTR* vector pJW1841 (Ashley et al. 2021) to generate pJW1934. The mScarlet cassette was then replaced with mNeonGreen (dpi) to generate pJW2229. The *mlt-11p (-2.8kb) mNeonGreen (dpi)-tbb-2 3’UTR* fragment was PCR amplified from pJW2229 and Gibson cloned into *Sph*I-HF+*Spe*I-HF double digested RMCE integration vector pLF3FShC to produce pJW2337. This vector was integrated into NM5179 and the SEC was excised as previously described (Nonet 2020).

A pJW2361 *mNeonGreen(dpi)::3xFLAG::PEST-tbb-2 3’UTR* vector for SapTrap with ATG and GTA connectors was constructed by linearizing pJW2322 (Clancy et al. 2023) by PCR and Gibson cloning in PCR amplified *mNeonGreen (dpi)::3xFLAG* from pJW2172 and a *linker::PEST::tbb-2 3’UTR* from pJW1836 (Ashley et al. 2021). The pJW2286 *mlt-11p (-2.8 kb)* promoter for SapTrap cloning was previously described (Clancy et al. 2023). The *mlt-11* promoter fragment from Frand *et al*. (2005) and the *mlt-11p (-5.3 kb)* promoter fragment were PCR amplified from a fosmid containing the *mlt-11* gene and Gibson cloned into linearized pJW2286 to make pJW2451 and pJW2457, respectively. These *mlt-11* promoter plasmids were SapTrap cloned with pJW2361 and pNM4216 to generate insertion vectors for rapid RMCE (Schwartz and Jorgensen 2016; Nonet 2023). The remaining promoter reporters were constructed by SapTrap cloning and rapid RMCE (Schwartz and Jorgensen 2016; Nonet 2023). A pJW2365 *pes-10 minimal promoter::mNeonGreen(dpi):: 3xFLAG::PEST-tbb-2 3’UTR* vector for SapTrap with ATG and GTA connectors was constructed by linearizing pJW2361 and Gibson cloning in a *pes-10* minimal promoter amplified from pJW1947 (Ashley et al. 2021). We chose candidate *cis-regulatory* elements through NHR-23 ChIP-seq peaks called by the modENCODE bioinformatic analysis and additional areas of open chromatin identified through ATAC-seq (Gerstein et al. 2010; Serizay et al. 2020). The ATAC-seq and NHR-23 ChIP-seq peak DNA sequences were PCR amplified from pJW2337 or pJW2457 and Gibson cloned into pDONR221 with TGG and ATG connectors for SapTrap. These plasmids were then SapTrap cloned with pJW2365 and pNM4216 to generate insertion vectors for rapid RMCE (Schwartz and Jorgensen 2016; Nonet 2023).

For the *mlt-11* internal knock-in (JDW541), the mNeonGreen::3xFLAG cassette was inserted in an unstructured region of exon 7. This strain was created by injection of RNPs [700 ng/µl IDT Cas9, 115 ng/µl crRNA and 250 ng/µl IDT tracrRNA] and a dsDNA repair template (25-50 ng/ul) created by PCR amplification of a pJW2172 plasmid template into N2 animals (Paix et al. 2014; Paix et al. 2015)(Supplementary Table S1). PCR products were melted to boost editing efficiency, as previously described (Ghanta and Mello 2020). crRNAs used and repair template oligos for deletions are provided in Supplementary Table S1. F1 progeny were screened by mNeonGreen expression.

*mlt-11* promoter region deletion strains were created by injection of Cas9 ribonucleoprotein complexes (RNPs)(Paix et al. 2014; Paix et al. 2015) [700 ng/µl IDT Cas9, 115 ng/µl each crRNA and 250 ng/µl IDT tracrRNA], oligonucleotide repair template (110 ng/µl) and pSEM229 co-injection marker (25 ng/µl)(El Mouridi et al. 2020) for screening into N2 or JDW541. Where possible, we selected “GGNGG” crRNA targets as these have been the most robust in our hand and support efficient editing (Farboud and Meyer 2015). F1s expressing the co-injection marker were isolated to lay eggs and screened by PCR for the deletion. Genotyping primers are provided in Supplementary Table S1. All deletions were confirmed by Sanger sequencing.

*mlt-11* coding sequence deletion strains were created by injection of Cas9 ribonucleoprotein complexes (RNPs)(Paix et al. 2014; Paix et al. 2015) [(700 ng/µl IDT Cas9, 115 ng/µl each crRNA and 250 ng/µl IDT tracrRNA) or (250 ng/ul IDT Cas9, 20 ng/ul each crRNA and 40 ng/ul IDT tracrRNA)], oligonucleotide repair template (110 ng/µl) and pSEM229 co-injection marker (25 ng/µl)(El Mouridi et al. 2020) for screening into strain EG7968. crRNA target selection, F1 progeny screening, isolation and genotyping were similar to our promoter region deletion workflow. We balanced the homozygous lethal mutations genetically by crossing to a strain with a *myo-2p::GFP::unc-54 3’UTR* cassette inserted into F46B3.7, a gene roughly 40 kb away. Homozygous viable lines were not crossed to the balancing strain. Genotyping primers are provided in Supplementary Table S1.

### Imaging

Synchronized animals were collected by either picking or washing off plates. For washing, 1000 µl of M9 + 2% gelatin was added to the plate or well, agitated to suspend animals in M9+gelatin, and then transferred to a 1.5 ml tube. Animals were spun at 700xg for 1 min. The media was then aspirated off, and animals were resuspended in 500µl M9 + 2% gelatin with 5 mM levamisole. 12 µl of animals in M9 +gel with levamisole solution were placed on slides with a 2% agarose pad and secured with a coverslip. For picking, animals were transferred to a 10 µl drop of M9+5 mM levamisole on a 2% agarose pad on a slide and secured with a coverslip. Images were acquired using a Plan-Apochromat 40x/1.3 Oil DIC lens or a Plan-Apochromat 63x/1.4 Oil DIC lens on an AxioImager M2 microscope (Carl Zeiss Microscopy, LLC) equipped with a Colibri 7 LED light source and an Axiocam 506 mono camera. Acquired images were processed through Fiji software (version: 2.0.0- rc-69/1.52p). For direct comparisons within a figure, we set the exposure conditions to avoid pixel saturation of the brightest sample and kept equivalent exposure for imaging of the other samples.

### RNAi Knockdown

RNA interference experiments were performed as in Johnson *et al*. (2023). Control RNAi used an empty L4440. The *mlt-11* RNAi vector was streaked from the Ahringer library (Kamath et al. 2003).

### Hoechst staining

Hoechst 33258 staining was performed as described previously (Moribe et al. 2004), except that we used 10 μg/ml of Hoechst 33258 as previously described (Ward et al. 2014). Two biological replicates were performed examining 50 animals per experiment. Representative images were taken with equivalent exposures using a 63× Oil DIC lens, as described in the imaging section.

## Data availability

A full description of all oligonucleotides, plasmids, transgenes, and *C. elegans* strains created and used in this article is in the set of supplemental tables. The authors affirm that all data necessary for confirming the conclusions of the article are present within the article, figures, tables, and Supplementary material. Plasmid sequences are provided in File S1, knock-in sequences from genome editing are provided in File S2. Any additional information is provided upon request.

## Acknowledgements

We would like to thank David Fay, Meera Sundaram, Andrew Chisholm, and Nathalie Pujol for helpful conversations. Andrew Chisholm and Nathalie Pujol provided invaluable feedback on the manuscript. We thank Krista Myles, Patricia Bliatout, Zoe Johnson, Javier Hernandez Lopez, Zoie Reyna, Emma Cadena, Valarie Hallin, and Olivia Vedar for research support. We thank Andrew Chisholm, Michael Nonet, Xiaochen Wang and Meera Sundaram for strains. Some strains were provided by the *Caenorhabditis* Genetics Center, which is funded by the NIH Office of Research Infrastructure Programs [P40 OD010440].

## Author Contributions

Conceptualization: J.M.R, J.D.W.

Methodology: J.M.R, A.T, A.J., A.A.V., V.P., K.D., J.C.C, M.T.L., A.L., J.D.W.

Validation: J.M.R, A.T, A.J., A.A.V., V.P., K.D., J.C.C, M.T.L., A.L., J.D.W.

Formal analysis: J.M.R, A.T, A.J., A.A.V., V.P., K.D., J.C.C, M.T.L.,A.L., J.D.W.

Resources: J.M.R, J.D.W.

Data curation: J.M.R, J.D.W.

Writing - original draft: J.M.R, J.D.W.

Writing - review & editing: J.M.R, A.T, A.J., A.A.V., V.P., K.D., J.C.C, M.T.L.,A.L., J.D.W.

Supervision: J.M.R, J.D.W.

Project administration: J.D.W.

Funding acquisition: J.D.W.

## Figure legends

**Supplementary Figure S1. Cuticle collagen and alae structure are disrupted in *mlt-11* promoter element deletion strains.** (A) ROL-6::mNG::3xFLAG localization in wild-type (control) and promoter deletion strains. (B) BLI-1::mNG::3xFLAG localization in wild-type (control) and promoter deletion strains. (C) DIC images of alae from wild-type (control), promoter deletion (peaks 3+4Δ) and knockdown (*mlt-11(RNAi))* L4+1 day worms. Images represent a minimum of 30 worms over multiple observations. Scale bars: 10 µm in A-B, 5 µm in C.

**Figure S2. MLT-11 Kunitz domain alignment.** Alignment of the ten MLT-11 Kunitz domains to Bovine Pancreatic Trypsin Inhibitor (BPTI). Positions of the six cysteine residues critical for the structure of each Kunitz domain are indicated above. Blue shading indicates conserved sequences and the histogram at the bottom depicts the degree of conservation with a consensus sequence listed below.

**Figure S3. Alignment of MLT-11 homologs.** MLT-11 homologs from the indicated nematode species were aligned using Clustal Omega. The length in amino acids of each homolog follows the species and homolog name. To the left and right of the alignment are amino acid positions of the end residues for each protein. Blue shading indicates conserved sequences and the histogram at the bottom depicts the degree of conservation with a consensus sequence listed below. The positions of the *C. elegans* MLT-11 signal sequence, potential furin cleavage site, thyroglobulin domain, three lustrin, and ten Kunitz domains are indicated. We note that we cut off the extended *P. pacificus* C-terminus (amino acids 2626-3742) since no sequence aligned to it as all proteins terminated at the *C. elegans* MLT-11 stop codon. No predicted motifs are found in the *P. pacificus* C-terminus.

**Figure S4. Embryonic inviability in *mlt-11* abrogated worms.** The percentage of unhatched embryos from worms with either a *mlt-11* whole gene deletion or deletion of specific domains. Δ/+ worms are homozygous inviable and so were balanced and maintained as heterozygotes. These heterozygotes were plated as adults and allowed to lay eggs for 2 hours, then removed. The resulting embryos were allowed to develop for 48 hours and then scored for hatching. Worms were scored over two biological replicates.

